# Active Learning of Cortical Connectivity from Two-Photon Imaging Data

**DOI:** 10.1101/268599

**Authors:** Martín Bertrán, Natalia Martínez, Ye Wang, David Dunson, Guillermo Sapiro, Dario Ringach

**Affiliations:** Electrical and Computer Engineering, Duke University, Durham, North Carolina, USA; Statistical Science Program, Duke University, Durham, North Carolina, USA; BME, CS, and Math, Duke University, Durham, North Carolina, USA; Neurobiology and Psychology, Jules Stein Eye Institute, Biomedical Engineering Program, David Geffen School of Medicine, University of California, Los Angeles, California, USA

## Abstract

Understanding how groups of neurons interact within a network is a fundamental question in system neuroscience. Instead of passively observing the ongoing activity of a network, we can typically perturb its activity, either by external sensory stimulation or directly via techniques such as two-photon optogenetics. A natural question is how to use such perturbations to identify the connectivity of the network efficiently. Here we introduce a method to infer sparse connectivity graphs from *in-vivo*, two-photon imaging of population activity in response to external stimuli. A novel aspect of the work is the introduction of a recommended distribution, incrementally learned from the data, to optimally refine the inferred network.. Unlike existing system identification techniques, this “active learning” method automatically focuses its attention on key undiscovered areas of the network, instead of targeting global uncertainty indicators like parameter variance. We show how active learning leads to faster inference while, at the same time, provides confidence intervals for the network parameters. We present simulations on artificial small-world networks to validate the methods and apply the method to real data. Analysis of frequency of motifs recovered show that cortical networks are consistent with a small-world topology model.

## Introduction

A fundamental question of system neuroscience is how large groups of neurons interact, within a network to perform computations that go beyond the individual ability of each one. One hypothesis is that the emergent behavior in neural networks results from their organization into a hierarchy of modular sub-networks, or motifs, each performing simpler computations than the network as a whole [1].

To test this hypothesis and to understand brain networks in general we need to develop methods that can reliably measure network connectivity, detect recurring motifs, elucidate the computations they perform, and understand how these smaller modules are combined into larger networks capable of performing increasingly complex computations.

Here we focus on the first of these problems, which is a pre-requisite to the rest: the identification of network connectivity from *in-vivo*, two-photon imaging data. Advances in two-photon imaging are giving us the first look at how large ensembles of neurons behave *in-vivo* during complex behavioral tasks [2–4]. Developing methods capable of analyzing the connectivity between a large number neurons, from noisy, stochastic activations, and limited recording time, is a significant challenge.

It is often the case that we can probe the networks under study, instead of merely observing their ongoing activity. For example, in studying visual cortex we can select a specific visual stimulus [5–8], or we can stimulate individual neurons directly via two-photon optogenetics [9, 10]. This active observation has been shown critical for system identification beyond brain networks, and is the direction here pursued.

We introduce a method that infers a sparse connectivity graph from available simultaneous recording of external stimuli and individual neural spiking rates, and recommends the future distribution of external stimuli to apply in order to optimally refine the inferred network. We show how such iterative “active learning” leads to faster inference while at the same time providing confidence measurements for the computed network components. The proposed decision-making approach takes into account information we may already have about network connectivity, the stochastic nature of the neural responses, and the uncertainty of connection weights.

It is important to note that the inferred connectivity graph implies functional, but not necessarily anatomical, connectivity. The recovered functional connectivity must be interpreted carefully when the recorded neural population is only a subset of a larger population. A directed connection might be caused, for example, by a common (but unobserved) driving factor, or hidden intermediate nodes. Nonetheless, common driving factors associated with stimulus modulations are partially accounted for by including stimulus information, this effect has also been discussed in [11]

Our framework consists of modelling the neuron spike trains with a Poisson Generalized Linear Model (GLM) [12], using past spike trains and applied stimuli as regressors. This type of point process model on neural spiking activity was popularized by [13]. The coefficients of these regressors in the GLM make the edge weights of the directed network. A variable selection approach is used to make the network sparse (set most edges to zero). Active learning is then used to decide, at any given point in time, the next sets of stimuli that allow for optimal inference of the network connectivity. Fig 1 shows a visual representation of the proposed framework.

**Fig 1.**
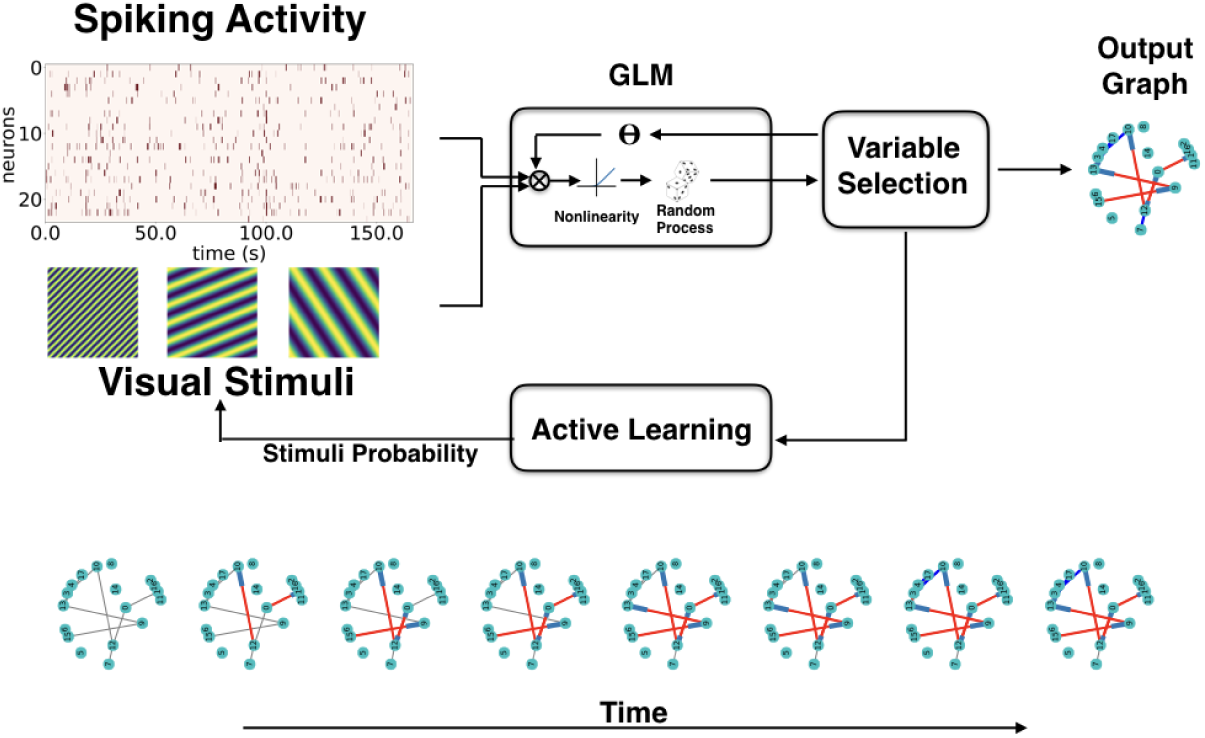
Recordings of spiking activity of a neuron population and the presented visual stimuli are fed into a GLM. The GLM and Variable Selection blocks work in tandem to decide which connections are relevant for explaining the system’s behaviour (the data) and building the directed connectivity graph (network). The active learning component analyzes the data obtained so far to optimize the visual stimuli to be presented for the next step of data acquisition, this is done to reduce graph uncertainty. This process is iteratively repeated. The bottom row shows how the network is gradually reconstructed as a function of acquired samples. Gray edges represent yet undiscovered edges present in the network, while red and blue edges represent discovered excitatory and inhibitory edges respectively.

While the proposed framework is general, we illustrate it by applying it to two-photon imaging data from mouse primary visual cortex. We also validate the method’s effectiveness on *in-silico* network simulations.

## Materials and methods

Following the brief description of the data acquisition, we then describe the foundation of the proposed active network inference framework. In doing so, we use terminology that will be relevant for the particular application at hand: the estimation of connectivity in neural networks. However, the framework is general enough to be applied in other contexts and using other modalities.

### Data acquisition

Animals: All procedures were approved by UCLA’s Office of Animal Research Oversight (the Institutional Animal Care and Use Committee), and were in accord with guidelines set by the US National Institutes of Health. The present study used data already collected for other studies. Thus, no new animal experiments were performed for the purposes of the present study. A detailed account of the experimental methods can be found elsewhere [14]. A brief description follows.

Imaging: Imaging of GCaMP6f expressed in primary visual cortex was performed using a resonant, two-photon microscope (Neurolabware, Los Angeles, CA) controlled by Scanbox acquisition software (Scanbox, Los Angeles, CA). The light source was a Coherent Chameleon Ultra II laser (Coherent Inc, Santa Clara, CA) running at 920nm. The objective was an x16 water immersion lens (Nikon, 0.8NA, 3mm working distance). The microscope frame rate was 15.6Hz (512 lines with a resonant mirror at 8kHz). Eye movements and pupil size were recorded via a Dalsa Genie M1280 camera (Teledyne Dalsa, Ontario, Canada) fitted with a 740 nm long-pass filter that looked at the eye indirectly through the reflection of an infrared-reflecting glass. Images were captured at an average depth of 260 µm.

Sequences of pseudo-random sinusoidal gratings [5, 15] and sparse noise stimuli were generated in real-time by a Processing sketch using OpenGL shaders (see http://processing.org). A detailed description is provided in [16]. The duration of the sequences was either 20 or 30 min, and gratings were updated 4 times a second on a screen refreshed at 60Hz. In a 20 min sequence, each combination of orientation and spatial frequency appeared at least 22 times on average.

In all experiments we used a BenQ XL2720Z screen which measured 60 cm by 34 cm and was viewed at 20 cm distance, subtending 112 x 80 degrees of visual angle. The screen was calibrated using a Photo-Research (Chatsworth, CA) PR-650 spectro-radiometer, and the result used to generate the appropriate gamma corrections for the red, green and blue components via an nVidia Quadro K4000 graphics card. The contrast of the stimulus was 80%. The center of the monitor was positioned with the center of the receptive field population for the eye contralateral to the cortical hemisphere under consideration. The location of the receptive fields were estimated by an automated process where localized, flickering checkerboards patches, appeared at randomized locations within the screen. This experiment was run at the beginning of each imaging session to ensure the centering of receptive fields on the monitor.

Image processing: The image processing pipeline was the same as described in detail elsewhere [14]. Briefly, calcium images were aligned to correct for motion artifacts. Following motion stabilization, we used a Matlab graphical user interface (GUI) tool developed in our laboratory to define regions of interest corresponding to putative cell bodies manually. Following segmentation, we extracted signals by computing the mean of the calcium fluorescence within each region of interest and discounting the signals from the nearby neuropil. Spikes were then estimated using the algorithm described in [17], available at https://github.com/darioringach/Vanilla. The present results are based on the inferred spiking activity.

### Generalized linear models: Poisson point process

Let 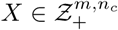 and 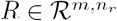 be two sets of random matrices called the target variables and regressor variables respectively. Each column *c* of the *X* matrix, denoted 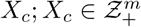 contains the spike train series of neuron *c*. Similarly, each column *r* of the *R* matrix, denoted *R*_*r*_; *R*_*r*_ *∈ ℛ*^*m*^, is a time series that can be either the past spiking activity of an observed neuron or the history of one of the presented external stimuli.

We say that the set *X* is a Poisson Point Process if the conditional probability distribution (CPD) of each column *X*_*c*_ is of the form

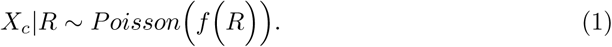

where f is a non-negative function, ∀*c* = 1*, …, n*_*c*_.

A Poisson Generalized linear model (GLM) is a special case of a general Poisson point process where

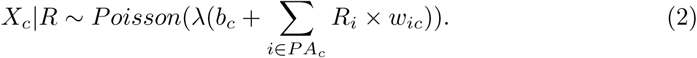

*λ* : **R** → **R**^+^ is a predefined nonlinearity, *w*_*ic*_ are the influence weights, *b*_*c*_ is the bias weight, and *PA*_*c*_ is the parent set of *X*_*c*_, and is the subset of the regressors *R* that carry information on the behaviour of neuron *c*. In our case *PA*_*c*_ consists of visual stimuli and past neuron activity that directly affect the spiking rate of neuron *c*. The parameter *w*_*ic*_ represents the magnitude of that influence.

The overall goal of network learning is finding the set (*PA*_*c*_*, w*_*ic*_), for each neuron *c*, that best represents the recorded data (stimuli and neural activity).

In the following sections we will present the model in more detail and the chosen algorithm for selecting the relevant regressors *PA*_*c*_ and influence weights *w*_*ic*_. Then we will introduce an experimental design method to select the stimuli that are more relevant in discovering the structure of the network. We want to emphasize that we are especially interested in correctly inferring the regressor set *PA*_*c*_, this can be seen as a binary classification problem, since a given regressor either belongs or does not belong on *PA*_*c*_ for any given neuron *c*. This will directly translate into the decision of which sets of stimuli are relevant. This decision will be based on improving the *PA*_*c*_ classification performance.

### Model description

The system under study consists of a set of *n*_*c*_ neurons {*C*} and *n*_*s*_ stimuli or external inputs {*S*}, that can be used to perturb the neuron’s activity.

To differentiate the influence of spiking activity and visual stimuli we will split the set of regressors *R* explicitly into spike activity *X*_*j*_(*t*)*, j* = 1*,‥, n*_*c*_, and stimuli activity *I*_*i*_(*t*)*, i* = 1*,‥, n*_*s*_.

To incorporate information about the past observations we further redefine the regressors as 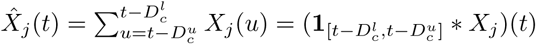, *j* = 1, …, *n_c_* and 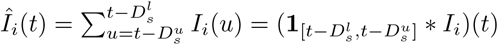, *i* = 1, …, *n_s_*, which are the convolution of past spiking activity (*X*_*j*_*, j* = 1,‥,*n*_*c*_) and past stimuli activity (*I*_*i*_*, i* = 1,‥,*n*_*s*_) with the boxcar influence function up to delays 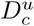 and 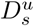 respectively.

To model the spiking train of neuron *c*, we also define the sets of relevant regressors of neuron *c*: 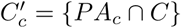 and 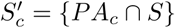.

Under these conditions, the spiking train of neuron *c* can be modeled as

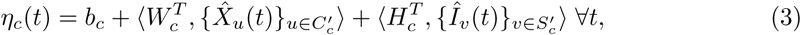

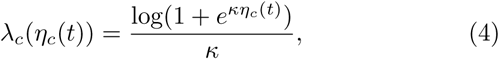

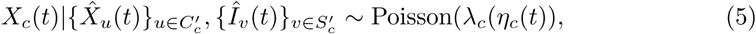

where *<. >* denotes inner product. The parameters of interest are:

- 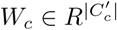 Edge weights between parent neurons in 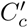 and *c*. These weights collectively represent the inter-neuron connectivity matrix

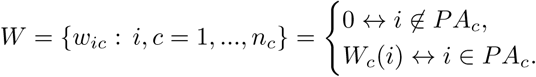
- 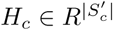 Edge weights between parent stimuli in 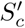 and *c*. These weights collectively represent the direct stimuli response matrix

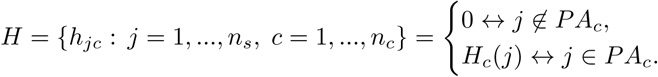
- *b*_*c*_ ∈ *R*: Bias of neuron *c*. This number encodes the base spiking rate of neuron *c* independently of the state of the other regressors.

Note that the model is time homogeneous, i.e., *W*_*c*_, *H*_*c*_ and *b*_*c*_ do not depend on time. The motivation for using the nonlinearity function Eq. (4) instead of the canonical exponential nonlinearity is that Eq. (4) is approximately exponential for low values of *η*_*c*_, but saturates to a linear dependency for large *η*_*c*_. This allows the model to capture strong inhibitions while at the same time ensuring excitations in the model do not grow out of control. The nonlinearity proposed in Eq. (4) depends on a static calibration constant *κ*, we provide additional insight into what this parameter does in the context of model selection in the Appendix.

### Parameter estimation

For each observed neuron *c* we want to find the regressor sets 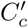 and 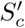 that best explain the data without over-fitting.

We first define the likelihood function of the proposed model,

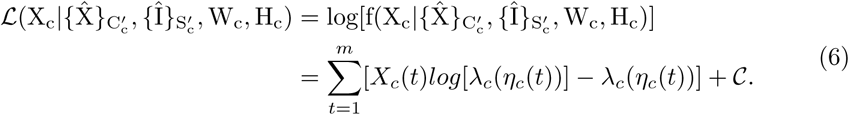

where 𝒞 is an additive constant A good regressor should provide a significant improvement in the model likelihood, and should have a tight confidence interval around its estimated edge weight. These notions are formalized using the Bayesian Information Criterion (BIC) [18] and the Wald test [19] respectively. A derivation of both in the context of our model is provided below.

To estimate the values for a set of parameters *θ* = {*W*_*c*_*, H*_*c*_*, b*_*c*_} we utilize the standard Maximum Likelihood Estimation (MLE) framework. The MLE estimate is denoted as 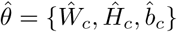 and is obtained as the solution of

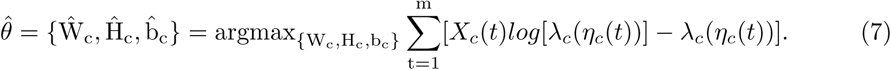

Furthermore, the parameter obtained from this estimation asymptotically follows a Normal distribution around the true value *θ*_0_ [20]. Under further regularity conditions [21], the variance of the estimator can be computed as shown in Eq. (8) (this will form the theoretical basis for the notion of tight confidence intervals),

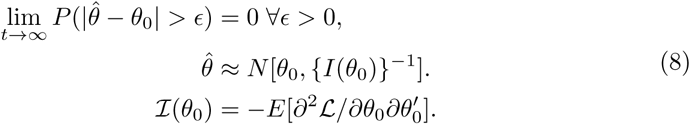

The quantity *ℐ*(*θ*_0_) is the Fisher information matrix [21]. Since we do not have access to the true parameter *θ*_0_ we use the observed Fisher information 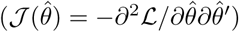 as an approximation. The observed Fisher information is sometimes referred to as the Hessian of the negative log-likelihood. The quantity ℐ{(*θ*_0_)}^-1^ is a lower bound for the variance of any unbiased estimator, as stated by the Cramér-Rao lower bound [22, 23]. The observed Fisher information of the model can be found on Eq. (29) in the Appendix.

It is important to note that, in general, the Fisher information matrix is singular in the under-sampled regime (i.e., when there are more parameters than observations). This issue is later circumvented by the regressor selection process described below. This is accomplished naturally since the largest observed variance is bounded from below with a number that grows arbitrarily large for ill-conditioned Fisher Information matrices. A more detailed explanation is given in the Appendix.

This Gaussian assumption in the non-asymptotic regime is partly justified by the fact that the log-likelihood function is concave with respect to the model parameters, with a single, well defined global optimum. This property is also shared by the Gaussian distribution. From these observations we realize that both the finite sample and the asymptotic parameter distributions are log concave and have a well defined global optimum. This makes the Gaussian approximation in the finite sample case a more reasonable approximation. This approximation has also been used in [24] for similar reasons

The MLE estimators combined with the observed Fisher information provide a confidence interval for each parameter of interest. We start from the null hypothesis *H*_0_ that the edge is irrelevant (zero value), and use the Wald test [19], to accept or reject this. The probability of parameter *θ*_*j*_ belonging to the null hypothesis can be computed as

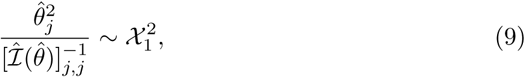

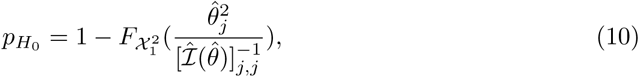

where 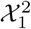 is the chi-square distribution with one degree of freedom, 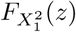 is the cumulative distribution function 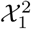 evaluated at *z*, and *pH*_*0*_ is the p-value associated with the null hypothesis.

At this point we have derived a measure of the probability of the parameter being different from zero.

To measure how informative a given regressor is we use the Bayesian information criteria (BIC). This quantity decreases with a higher likelihood 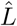 and increases with the number of parameters currently used in the model 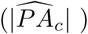, and the number of observations (*m*):

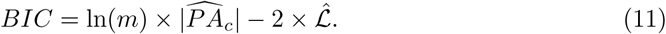

When comparing two models, the model with the lowest BIC value is preferred. The quantity 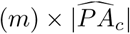 penalizes model complexity, reducing the number of noisy weak connections.

This quantity is introduced to favor sparsely connected networks, since edges that have a low overall effect on the predicted likelihood of the model are ignored in favor of model sparsity.

For each observed neuron *c*, our objective is finding the parent set estimate 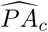 that minimizes the BIC, subject to a p-value restriction *γ* which forces the selected regressors to have a tight confidence interval:

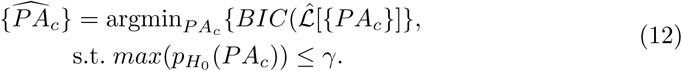

This combined approach looks for a sparse model representation able to capture most of the information present in the observed spike trains using a series of parameters (edge weights) with well-defined values, with tight uncertainty intervals.

In the following section we will briefly explain how we approach this minimization.

### Selecting relevant regressors

The optimization problem presented in Eq. (12) can be stated concisely as the problem of finding the set of regressors 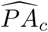 that yields the best BIC score (Eq. 11) subject to a p-value restriction. This is done to ensure that the regression model has good prediction capabilities and generalizes well to non observed data points.

The problem as stated in Eq. (12) is combinatorial in nature and cannot be directly optimized. There is a rich literature on model selection using various search strategies and evaluation criteria [25–31]. Usual model evaluation criteria include BIC and Akaike Information Criterion (AIC) [32] among others, while search algorithms include stochastic search variable selection [31], forward model selection, backward elimination, and stepwise methods in general, among others [26].

For this particular problem, we decided to use a greedy elastic-forward subset selection algorithm; an extensive overview of subset selection strategies can be found in [26]. In addition, we take several randomly selected subsets from the training dataset, each containing a fraction *ν* of available samples, and we evaluate the BIC performance and p-value restriction for regressor candidates across all bootstrapping subsets to find candidates that are consistently relevant across subsets. In our experiments the fraction *ν* is set to 0.7. We show the performance of varying the *ν* parameter on simulated data in the Appendix.

The algorithm starts from *PA*_*c*_ = Ø, and iteratively includes regressors that improve the median BIC score across the random subsets, as well as the BIC score over the full dataset, while satisfying the p-value constraint.Since this algorithm is rerun for every new batch of data, this effectively means that edges can be “removed” between successive data acquisitions The algorithm is described in detail in the Appendix. The results section compares the performance of this algorithm against the LASSO method for variable selection [33].

Now that we have described the model and algorithm, we proceed to describe the active learning strategy.

### Experimental design - Active learning

Our goal is to develop a method to select, at any time, the optimal action (or network perturbation) that is expected to yield the maximum information about its currently computed connectivity. For the purposes of this paper, our action set will consist of selecting which set of visual stimuli will be presented next.

We are interested in gathering samples from network connections (edges) that show a promising improvement in the likelihood of the model but have not yet been added to it, we refer to these edges as candidate edges. That means we want to improve the parent set estimate 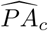*, c* = 1*, …, n*_*c*_, and edge weight estimates *w*_*ic*_*, i* = 1*, …, n*_*r*_*, c* = 1*, …, n*_*c*_, as in equations (12) and (7) respectively. The samples are collected by presenting stimuli that directly or indirectly generate activations in the parent nodes of the promising candidate edges.

To address this, we define a relevance score for each potential stimulus. This score considers the expected rate change of every regressor when presenting a given stimulus more frequently than the rest, weighted by the deviance statistic [34] of every edge associated with that regressor not currently present in the model. This in effect means that stimuli that directly trigger neurons associated with good candidate edges will be presented more often than other stimuli during the next intervention. The exact formulation is presented next.

### Defining a score for each stimuli

Define 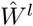, 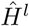 as the estimated adjacency and stimuli response matrices up to sample *m*_*l*_, where *l* is an iteration counter. Our objective is to obtain a probability distribution vector 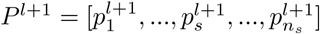 for presenting each stimulus *s* = 1*, …, n*_*s*_ at intervention *l* + 1.

Future stimuli sequences will be sampled from this distribution. We introduce this intermediary stimuli distribution probability instead of simply applying the stimulus with the highest expected change in utility for two reasons. Firstly, we can sample several consecutive stimuli from this distribution, this way, we can apply a small batch of stimuli before recomputing the optimum stimulus, reducing the computational costs. Secondly, this approach is less greedy than exclusively presenting the stimulus that has the highest expected change in utility. This can be beneficial in settings where the inferred model has high uncertainty.

We start by computing the expected firing rate change on every neuron *c* = 1*, …, n*_*c*_ of presenting each stimulus *s* = 1*, …, n*_*s*_ more frequently. For each stimulus *s*, we define a surrogate probability distribution vector associated with it: 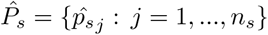, where stimulus *s* has the highest occurrence probability. Given the previous observations, we want to predict the expected change in spiking rate of each neuron *c* when preferentially applying stimulus *s* using 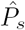, as opposed to the baseline where all stimulus are presented equally: 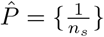. Formally the probability vector that favors stimulus *s* 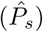 is defined as

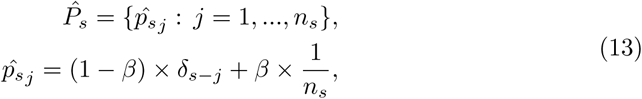

where 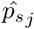 is the *j*-th element of the probability vector 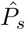 and corresponds to the probability of presenting stimulus *j* in the surrogate probability distribution vector associated with *s*, *β* is a smoothing constant that satisfies 0 ≤ *β* ≤ 1 and controls the overall probability of other stimuli appearing, and *n*_*s*_ is the number of available stimuli. We will define 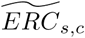 as the expected rate change of stimulus *s* on neuron *c*.

The estimated firing rate change on neuron *c* caused by stimulus *s* at iteration *l* + 1 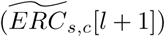 is presented in Eq. (14) next. This compares the expected firing rate of using a stimuli distribution 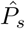 on neuron *c* when compared to using the baseline (uniform) distribution 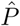. The quantity 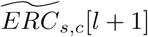 shows the rate increase of preferentially presenting stimuli *s* on the spiking rate of neuron *c* (*λ*_*c*_) according to our previous *m*_*l*_ observations,

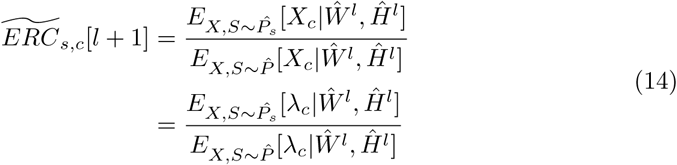

Next we compute the deviance statistic [34] for every possible outbound edge of neuron *c* that did not satisfy the parameter selection criteria in Eq. (12) and therefore 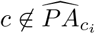 for some neuron *c*_*i*_. This is twice the log-likelihood difference (noted as 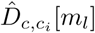) between the network model up to sample *m*_*l*_ (where 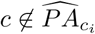) and a network model where *c* is included as a parent of *c*_*i*_:

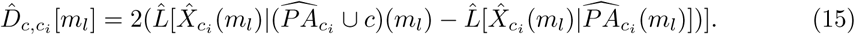

To compute the deviance statistic to any given edge from neuron *c* to neuron *c*_*i*_, we need to recompute the Maximum Likelihood (ML) estimate under the new regressor subset 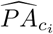. Fortunately, this ML estimate is fast to compute, the computation can be made faster by initializing the ML minimizer to the previously obtained regressor values.

Note that Eq. (15) is always non-positive, a highly negative value indicates a strong possibility of neuron *c* influencing neuron *c*_*i*_. By acquiring more samples from this interaction (samples where the candidate parent node is active), we can either disprove this notion, or gather enough evidence to add this edge into the regressor set (by satisfying the BIC and p-value criteria for adding an edge to the model).

We therefore define the score of stimulus *s* associated with inter neuron edges *W* as

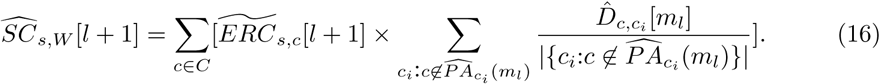

We have so far assigned a score that considers the deviance impact of preferentially applying a stimulus *s* on the inter neuron connectivity matrix *W*. It is important to note that the set 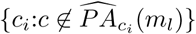 refers to the set of cells that up to sample *m*_*l*_ were not included as children of cell *c*, and therefore are considered as candidate edges for the purpose of the score. In this score, the first summation considers the impact of stimulus *s* over each cell *c* multiplied by the second summation, which is the mean log-likelihood difference over all the cells that were not considered as children of neuron *c* up to sample *m*_*l*_.

In a similar fashion, we also need to consider the effect of prioritizing any given stimulus *s* on the stimuli response matrix *H*. In this case, the expected rate change of prioritizing stimulus *s* over stimulus *s*_*i*_ is the quotient of the *s*_*i*_ entry of the stimuli probability distribution vector associated with *s* 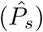 over the baseline (uniform) probability distribution vector 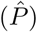,

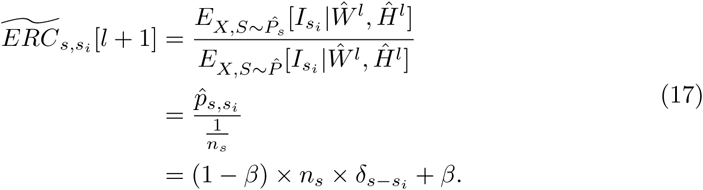

The deviance score of the output edges of *s* that did not satisfy the parameter selection criteria in Eq. (12) and the score of stimulus *s* associated with the stimuli response matrix *H* can be analogously defined:

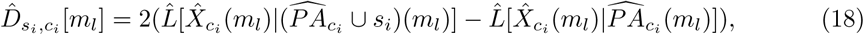

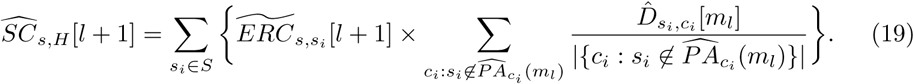

Finally, the combined score given to stimulus *s* is

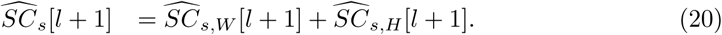

At this point we have a score for each stimulus *s* that is able to capture how informative this stimulus might be based on the expected rate change it has on edges that are not included in the model so far. The next step is mapping these scores into a probability vector. For that we first compute the z-score of each stimulus; this is done as a normalization step of the score values, and allows the detection of outlying stimuli. The z-scores are then converted into a probability vector with the use of the well known softmax function. To avoid giving unnecessarily small or large probabilities to any given stimulus, we truncate the computed z-score into the [–2, 2] range.

The formulation is as follows:

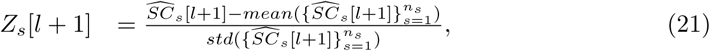

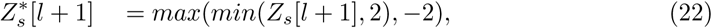

and the probability distribution vector for presenting each stimulus at intervention *l* + 1 ends up being:

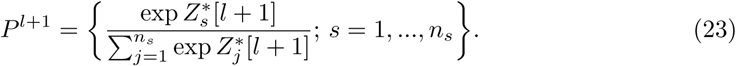

The use of the z-score as a normalization step allows the algorithm to dynamically pick up on the “relative quality” of the stimulation actions, and the truncation of the score provides a limit on how frequently or infrequently any given stimulus can be shown.

### Active learning

The active learning strategy consists of iteratively evaluating the current network model using Algorithm 1 in the Appendix, then using equations (20) and (23) to compute the stimuli distribution probabilities for the next time interval. The full algorithm is described in Algorithm 2 in the Appendix.

## Results

### Simulated data

In order to validate the method before applying it to real datasets, we generated a number of artificial datasets where the connectivity is known.

Network topology was simulated using the small-world Watts-Strogatz model [35]. This type of network architecture has been used to model functional cortical connectivity in cats and macaques [36, 37], and has been theorized to be of use in understanding human functional connectivity [38].

We defined two separate networks SW1CL and SW3CL, network SW1CL is a single small-world cluster network with 18 neurons, while network SW3CL has three separate small-world clusters, each cluster has 18 neurons. All clusters have an average connectivity degree of 0.03. The increase in spiking rate for neuron to neuron edges in network SW1CL were drawn from a normal distribution *N* (0.05, 0.005), the edge weights in the adjacency matrix were set accordingly. Similarly, the increase in spiking rate for neuron to neuron edges for the three clusters in network SW3CL were drawn from normal distributions *N* (0.075, 0.005), *N* (0.05, 0.005) and *N* (0.035, 0.005) respectively. Thirty percent of inter-neuron edges were made inhibitory.

These simulated networks were presented with 30 possible excitatory stimuli, most of which were designed to have no effect on the network. This was done to test that the active learning algorithm has the ability of navigating through confounders. The increase in spiking rate fro stimuli to neuron connections were drawn from the normal distribution *N* (0.10, 0.014). Fig. 2 shows the connectivity matrix *W* and stimuli response matrix *H* for networks SW1CL and SW3CL. Boxcar influence functions as defined for Eq. (3) were set to 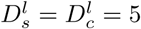 and 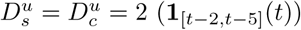. These values were selected so that the average spiking rate of the simulated neurons were similar to the ones obtained in real data.

**Fig 2.**
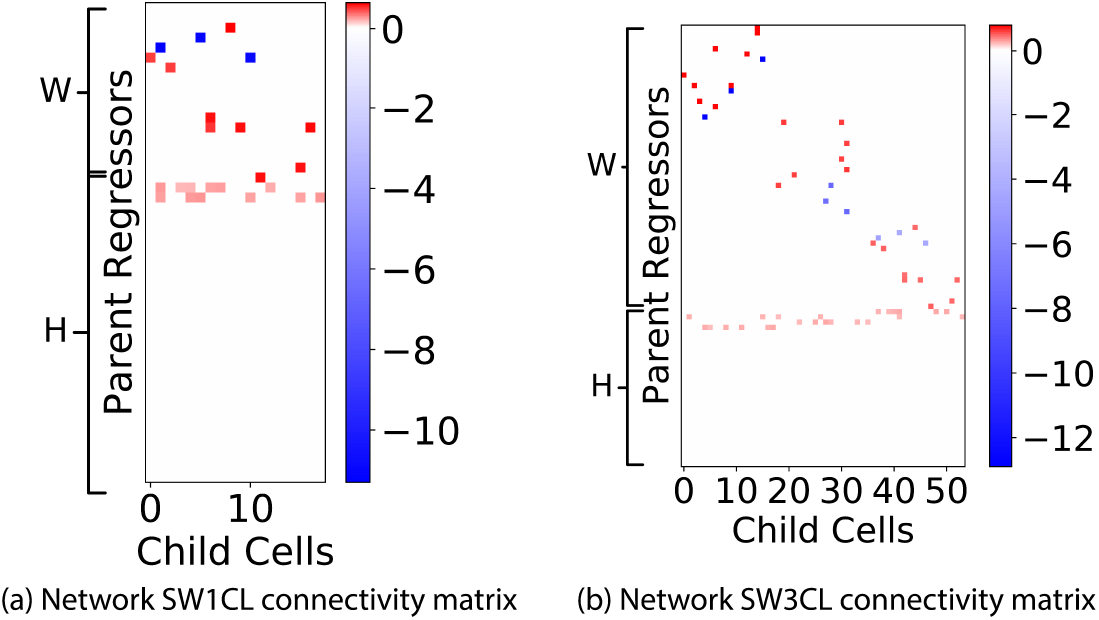
Adjacency matrices for networks SW1CL and SW3CL. Red entries in the adjacency matrices denote an excitatory relation between the regressor and the child neuron, while blue entries denote inhibitory connections. The block diagonal structure present in the *W* matrix for network SW3CL evidences the three cluster structure of the network. These connectivity matrices were computed to generate the required spiking rate change for model parameter *κ* = 10. On both networks, we can observe the large number of stimuli that have no effect on the network.

In the following sections we will present two experiments. The first experiment will show the performance difference in regressor selection when using the proposed Algorithm 1 compared against the Lasso method [33]. The second experiment will compare the performance in regressor selection when stimuli are chosen according to active learning Algorithm 2 versus random stimuli selection.

Algorithmic performance will be presented based on precision, recall, and *F*_1_ metrics:

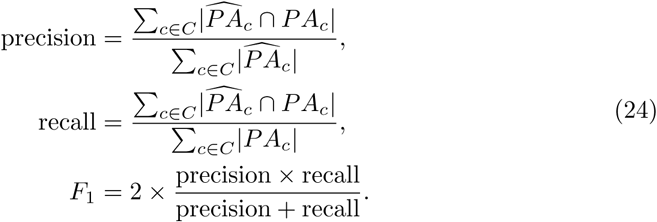

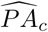 is again the recovered set of regressors, and *PA*_*c*_ is the true set of regressor edges for neuron *c*.

### Relevant regressors

We first checked the performance of the regressor selection methods on networks SW1CL and SW3CL. We compare the performance of the elastic-forward BIC selection Algorithm 1 with bootstrapping versus standard Lasso [33] regression using the pyglmnet implementation [39].

Stimuli were sampled uniformly with replacement, each stimulus was presented for 4 consecutive frames. Spiking trains for the simulations were sampled from a Poisson random process with a spiking rate corresponding to the ground truth model from Eq. (5).

The *l*_1_ regularization parameter for the Lasso method was selected using an oracle to provide the best possible *F*_1_ score. This method was selected for comparison because it is one of the most common approaches to variable selection. The modified log-likelihood function used for the Lasso method was:

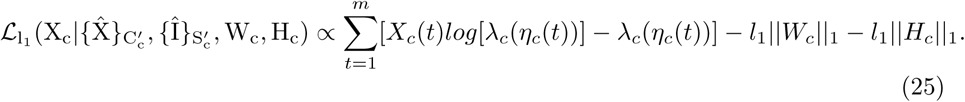

Fig. 3 shows the results; 10 independent trials were used to provide confidence intervals for the metrics.

**Fig 3.**
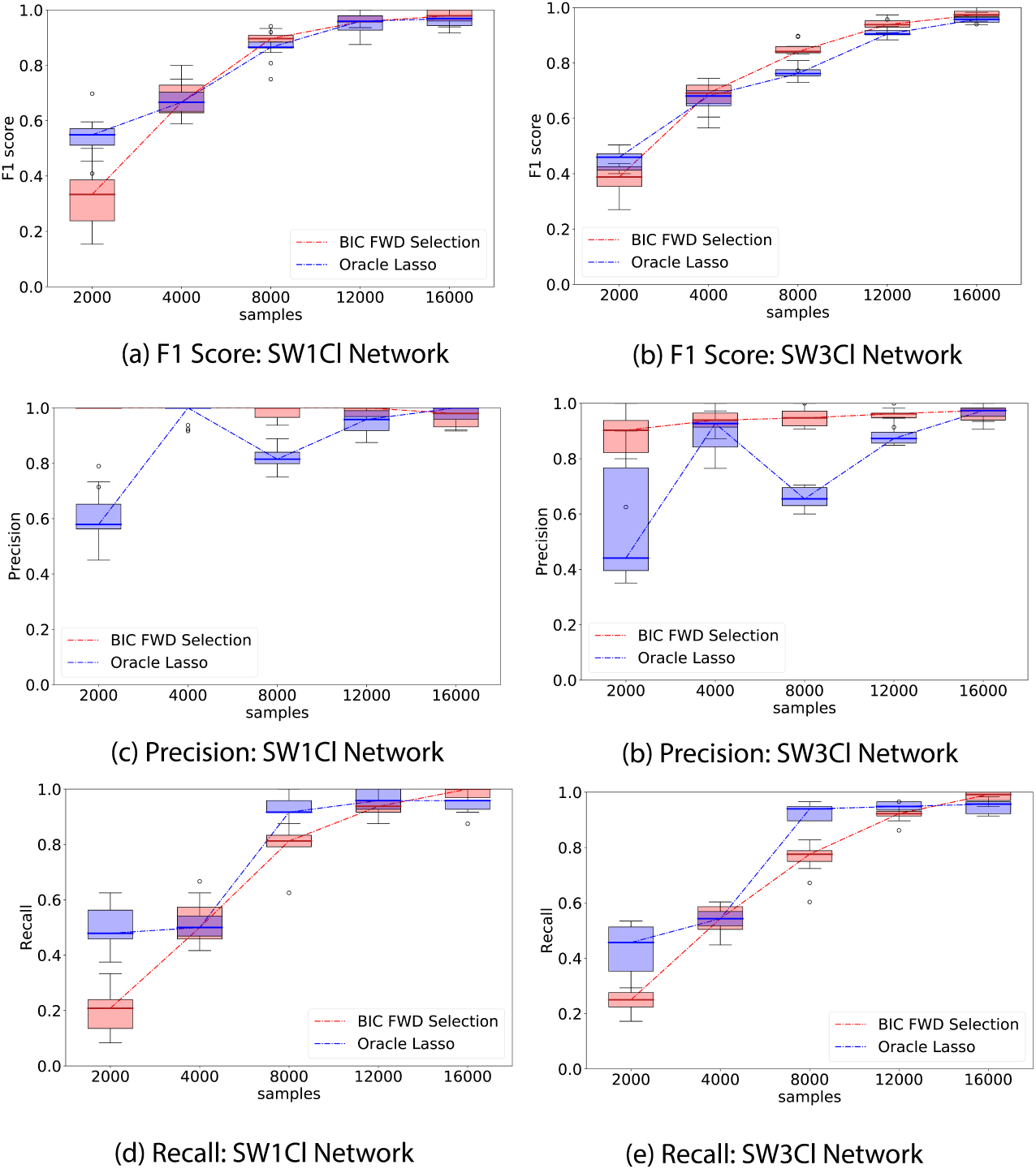
Whisker plot of performance indicators as a function of number of samples; elastic-forward BIC selection is shown in red, oracle lasso in blue. The whisker plot is obtained from 10 independent trials. External stimuli are drawn randomly from a uniform distribution; network SW1CL has a total of 24 non-zero parameters out 864 potential regressors, while network SW3CL has a total of 58 non-zero parameters out of 4536 potential regressors. The elastic-forward BIC selection outperforms the oracle lasso for larger sample sizes. This performance improvement is more noticeable in the SW3CL network, where edge weights are more diverse.

Fig. 3 shows that the elastic-forward BIC method described in Algorithm 1 outperforms lasso for larger sample sizes, even when the *l*_1_ regularization parameter is selected using an oracle. The improvement is more noticeable for network SW3Cl which has diverse edge weights. In all tested cases, the precision metric in edge recovery was better for elastic-forward BIC subset selection.

### Active learning: Stimuli selection

We now evaluate the performance of the proposed active learning method, Algorithm 2. We compare it against uniformly sampling from all 30 possible stimuli.

Both strategies start from the same initial 500 samples, and each intervention adds an additional 500 samples. At the beginning of each intervention step, we compute the best network estimate so far using Algorithm 1, and show the performance in recovering the set of regressor edges *PA*_*c*_ using the *F*_1_, precision and recall metrics. At this stage, the active learning strategy described in Algorithm 2 recomputes the stimuli probability distribution to apply for the following 500 samples. Active learning parameter *β* was set to *β* = 1*/*4.

Fig. 4 compares the performance of uniform (Random) stimuli sampling and active learning (AL) sampling, while Fig. 5 shows the performance difference only on the entries of the stimuli response matrix *H*. Experiments were repeated 10 times to provide confidence intervals for the metrics.

**Fig 4.**
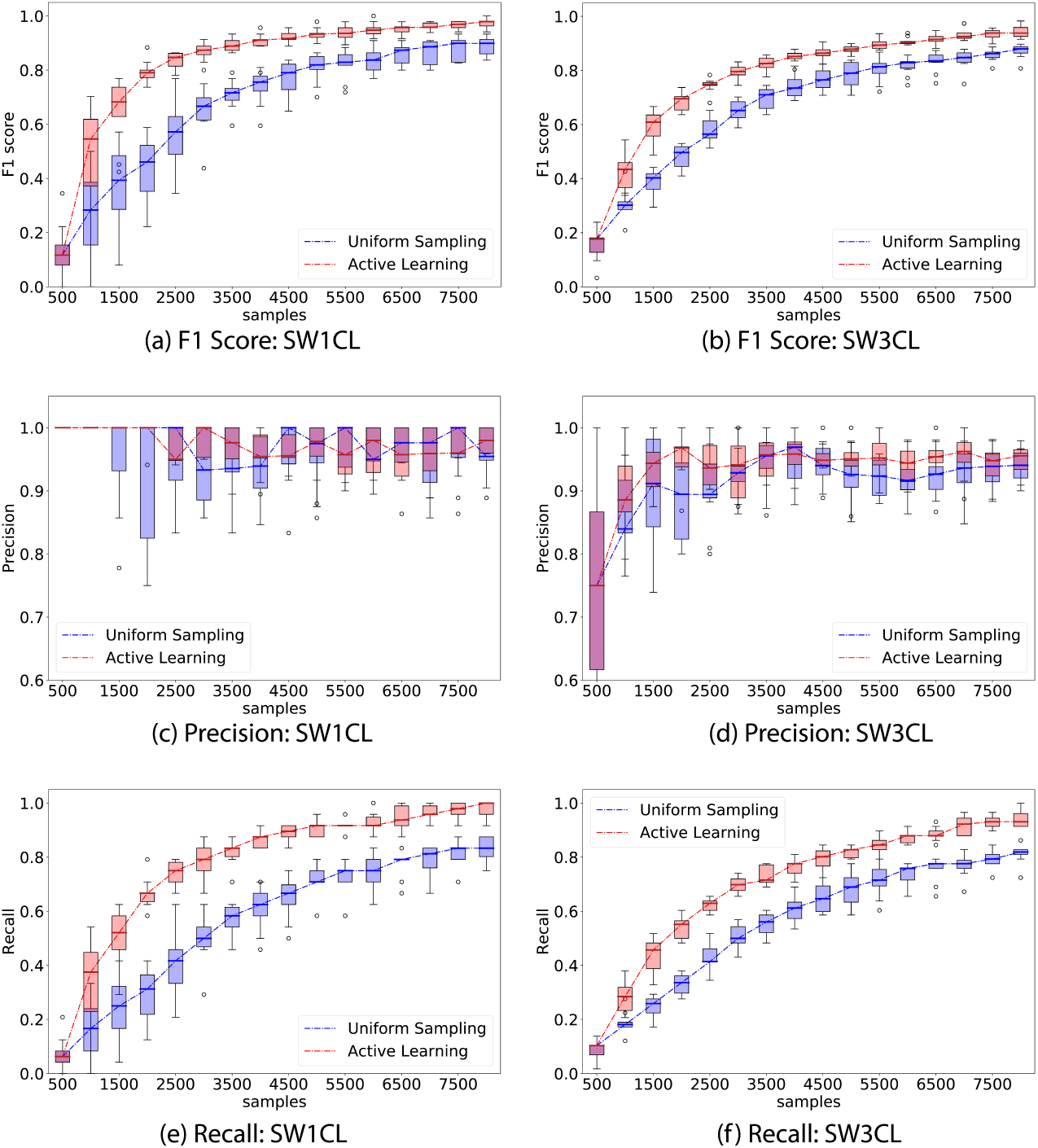
Comparison of performance between the proposed active learning (AL) method versus uniformly sampling (Random) from all stimuli. The experiment consisted of 500 sample interventions, with an initial 500 sample observation. Whisker plots are obtained from 10 independent trials. Left column show *F*_1_, precision, and recall performance on network SW1CL and right column on network SW3CL.

**Fig 5.**
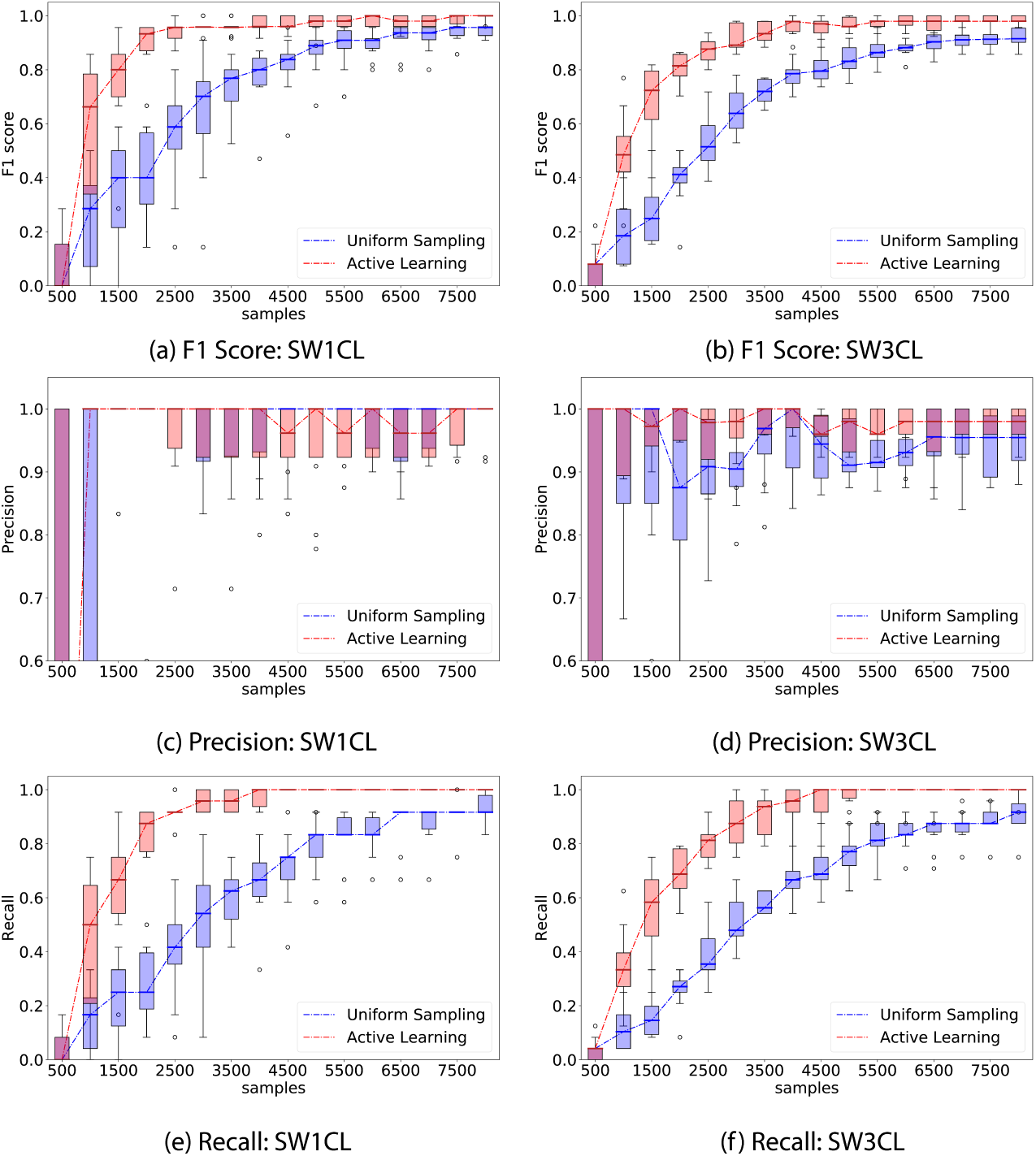
Comparison of performance between the proposed active learning (AL) method versus uniformly sampling (Random) from all stimuli only over the direct stimuli response matrices *H*. The experiment consisted of 500 sample interventions, with an initial 500 sample observation. Whisker plots are obtained from 10 independent trials. Left column shows *F*_1_, precision, and recall performance on network SW1CL and right on network SW3CL. On average, computing the active learning intervention step took 20*s* ± 9*s* on network SW1CL, and 98*s* ± 50*s* on network SW3CL.

Active learning outperforms random sampling by a large margin, the inter-quartile ranges for random sampling and active learning do not overlap over a significant sample count. As expected, both strategies converge for large sample sizes, but the process is sped-up by selecting the correct set of stimuli. The most noticeable performance difference is obtained when recovering the stimuli response edges *H*, since these are the edges we have direct influence on.

Fig. 6 shows a comparison between the elastic-forward BIC method described in Algorithm 1 and oracle lasso when applied to samples drawn from the recommended active learning distribution proposed in Algorithm 2. Both regressor selection methods perform similar under these conditions.

**Fig 6.**
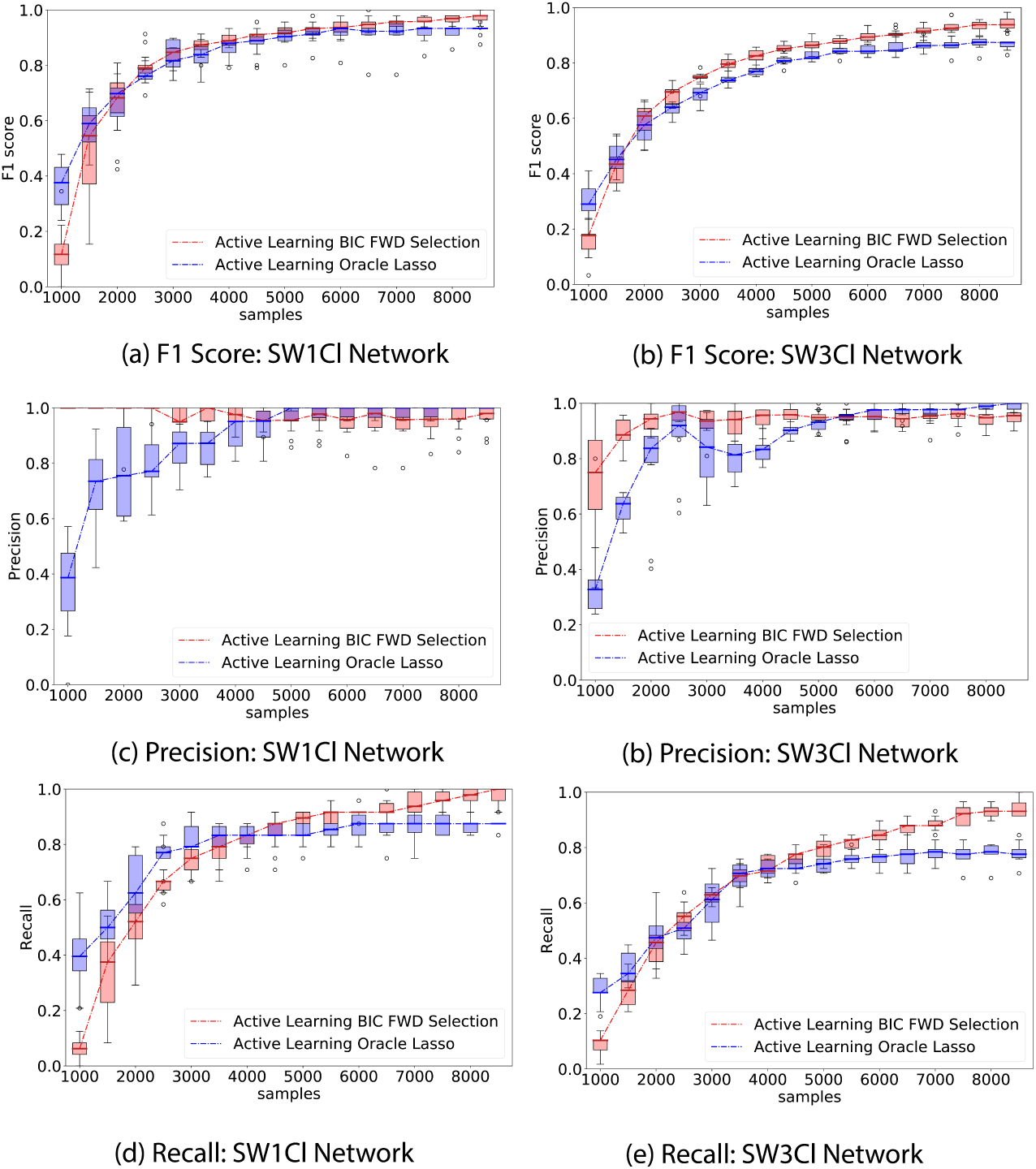
Whisker plot of performance indicators as a function of number of samples; elastic-forward BIC selection is shown in red, oracle lasso in blue. The whisker plot is obtained from 10 independent trials. External stimuli are drawn from the recommended Active Learning distribution; network SW1CL has a total of 24 non-zero parameters out 864 potential regressors, while network SW3CL has a total of 58 non-zero parameters out of 4536 potential regressors. The elastic-forward BIC selection still outperforms the oracle lasso for larger sample sizes. Both methods show a performance gain on the samples drawn from the Active Learning distribution when compared to their uniform sampling counterparts.

Fig. 7 shows a visual representation of the edge discovering process over network SW1CL using active learning versus random sampling. Fig. 8 shows the difference between ground truth and the active learning and random sampling estimates as a function of interventions. Here we can clearly see that *H* edges are quickly recovered using active learning, while *W* edges show a slower improvement that is network dependent.

**Fig 7.**
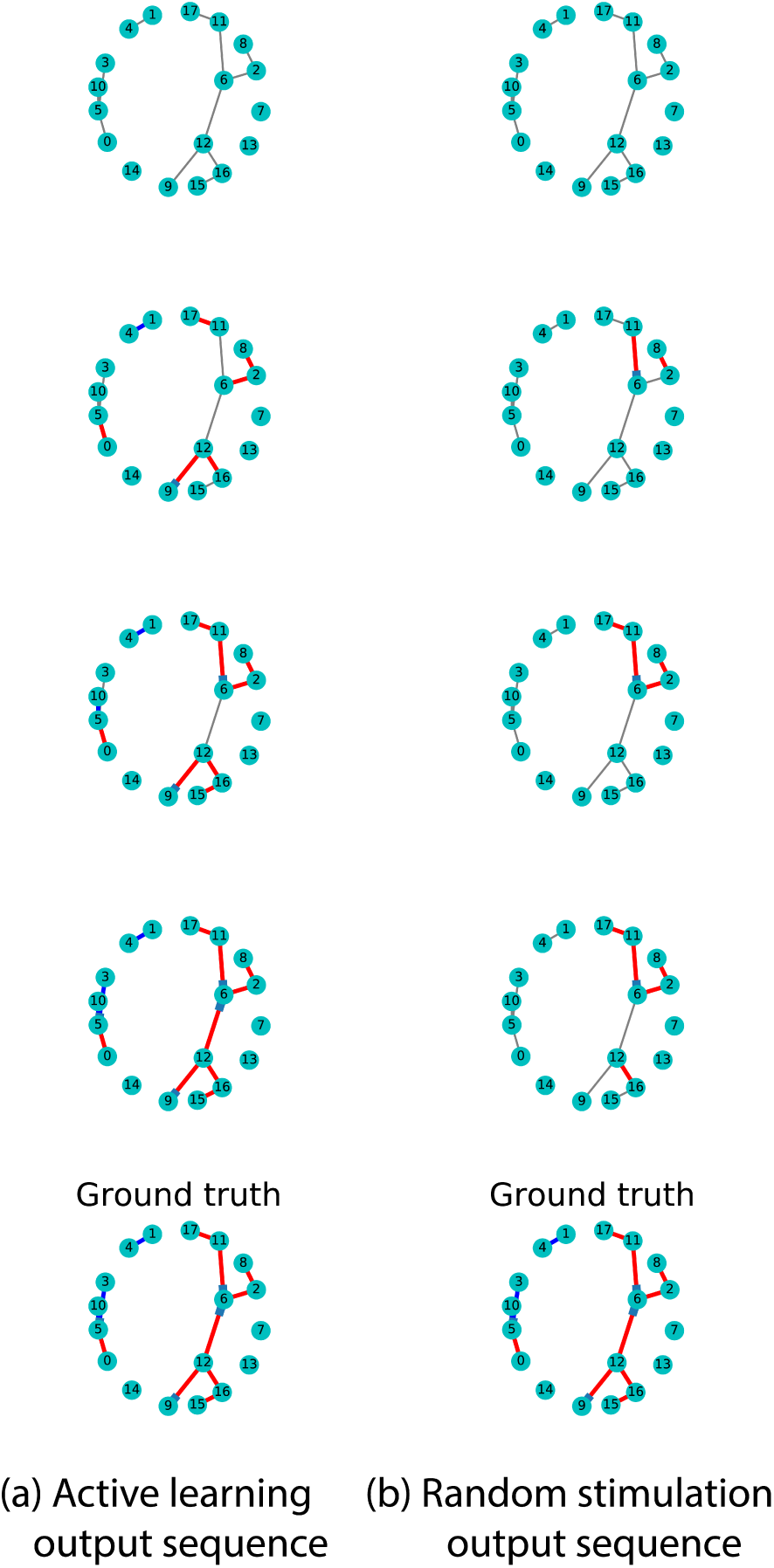
Comparison of the edge discovering process for network SW1CL using active learning versus random stimulation. Rows show inferred connections over a simulated cluster as a function of samples. Red and blue edges show correctly detected excitatory and inhibitory connections respectively, while grey edges show connections as not yet detected. Left column shows detected edges as a function of time when the active learning stimulation policy is used, right column shows the same cluster with a completely random stimulation policy. Rightmost cluster shows the ground truth. This example shows a clear advantage in edge recovery when using active learning compared to random stimulation.

**Fig 8.**
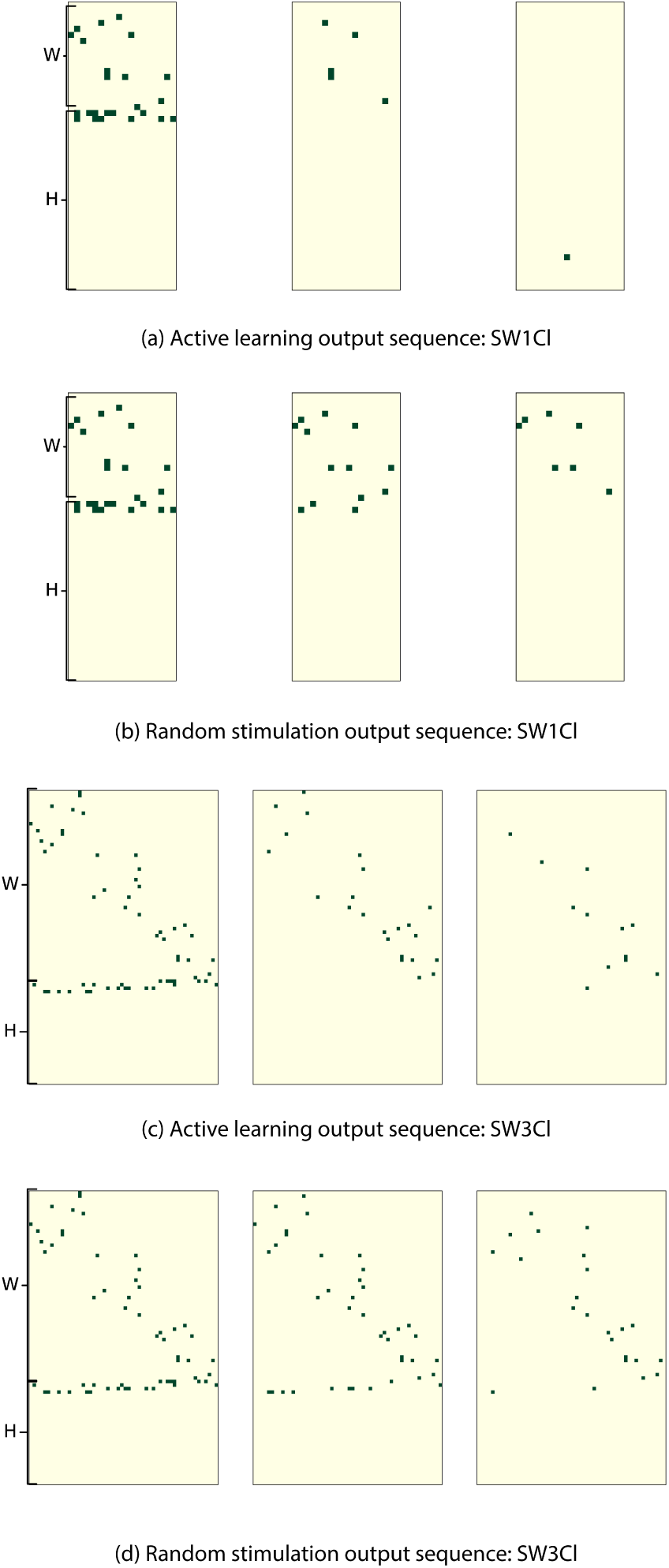
Comparison of the edge discovering process for networks SW1CL and SW3CL using active learning versus random stimulation. Rows show misclassified edges in the adjacency matrices *W* and *H* as a function of samples. Rows 1 and 3 show misclassified edges as a function of time when the active learning stimulation policy is used, while rows 2 and 4 show the same network probed with a random stimulation policy. The misclassified edge matrix under active learning quickly becomes sparse as the number of misclassifications goes to zero, random stimulation produces the same results but in a longer time frame.

### Real data

We worked with two datasets: lt3-000-002 and lt3-000-003, hereafter called datasets 1 and 2, containing a population of 57 and 63 neurons respectively. The presented visual stimuli consisted of sinusoidal gratings defined using Hartley basis functions of the form:

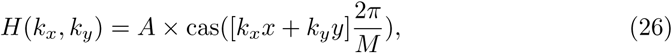

where the function *cas*(*x*) is *cos*(*x*) + *sin*(*x*), *A* = ±1, (*x, y*) are the pixel coordinates in the monitor, *M* is the image size, and *k*_*x*_*, k*_*y*_ = {-12*, …,* 12} are the frequency components. Parameters *A, k*_*x*_*, k*_*y*_ were uniformly sampled from all possible values, and each stimuli persisted for 4 frames.

To prevent excessive data fragmentation, the stimuli basis (*A, k*_*x*_*, k*_*y*_) was encoded into an (*r, Φ*) pair using Eq. (27) with the *r* and *Φ* parameters discretized into 7 values each,

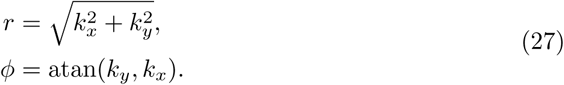

For both spike and stimuli regressors, we used **1**_[*t-*2*,t-*7]_(*t*) as the influence function as defined for Eq. (3) (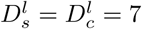 and 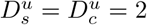).

7, 000 samples of each dataset were used for training, and 2000 samples were reserved for model validation. Models were computed under two conditions, the first model used all available regressors (full model), while the second model was restricted to self regression coefficients and direct stimuli to neuron connectivity (AR model).

We first show the predictive power of both models when evaluated over the validation samples, that is to say, we evaluate the log likelihood (Eq. (6)) of both models over the test samples.

We then utilize the model that we obtained from all samples in the dataset as a template for simulations (“ground truth”), and compare the simulated performance between the proposed active learning Algorithm 2 and random sampling.

We go on to show that observed neurons tend to respond more to low frequency stimuli. We then show the recovered adjacency matrix (networks) for both datasets, and spiking trains time series for neurons belonging to the largest cliques in the network.

### Recovered models

We now show the recovered connectivity matrices *W* and stimuli response matrices *H* for both datasets for the full model and the AR model.

The recovered inter-neuron connectivity matrix *W* for both datasets was 93% sparse. Both datasets also show a large number of stimuli connections in the *H* matrix corresponding to low spatial frequency stimulation values (*r*), and consistently selected the self regression coefficient as an important regressor. This was consistent for both regression models.

### Out of sample prediction power

To test the prediction power of the regressor model, we first evaluate the log likelihood of the full model and AR model over the test samples. We compute *η*_*c*_ and *λ*_*c*_ for each neuron on the test samples using the observed spike trains and visual stimuli as presented in equations (3) and (4). We then evaluate the log likelihood according to Eq. (6). The results are shown in Fig. 10.

**Fig 9.**
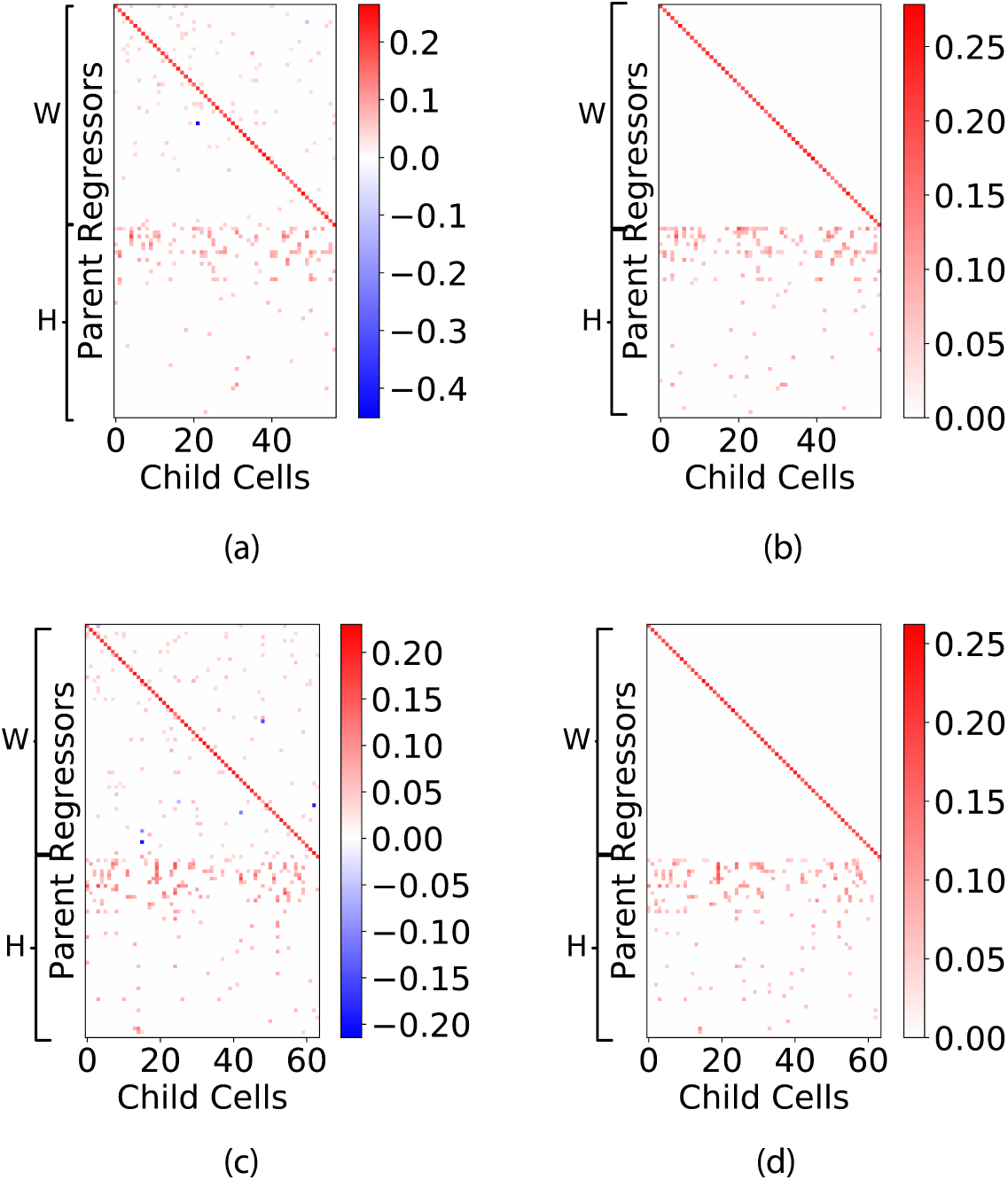
Recovered adjacency matrices for datasets 1 and 2. Top and bottom rows show the recovered adjacency matrices for datasets 1 and 2 respectively. Columns from left to right show the full model and the AR model respectively. Inhibitory connections are shown in blue, and excitatory connections are shown in red. We can observe that self regression coefficients are always added to the model. We also observe that the overall sparsity of the recovered network is consistent across datasets, and that the first few rows of the *H* matrices show a heavy concentration of excitatory connections. These rows correspond to low spatial frequency (*r*) values in the Hartley basis functions (Eq.(26)).

**Fig 10.**
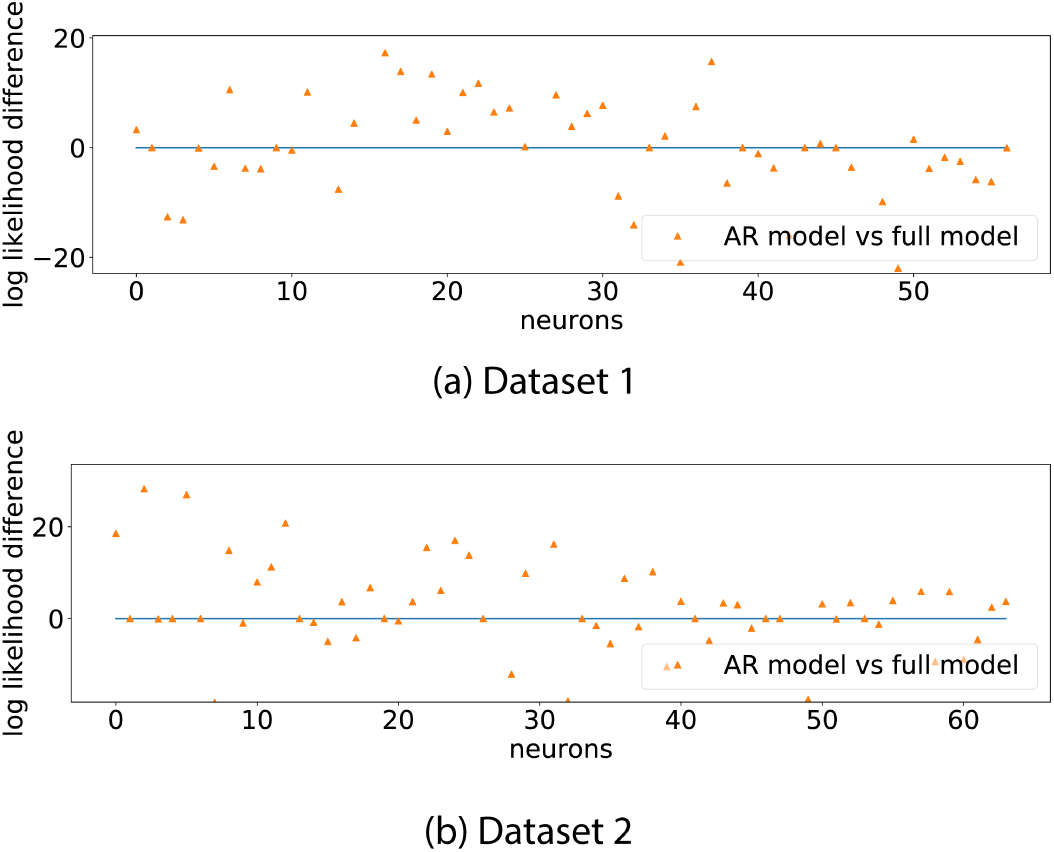
Log likelihood difference of full model and AR model over the test samples. The graph shows that, on average, spiking rate predictions are better on the full model than on the auto-regressive model. This grounds the idea of neuron to neuron interaction as being predictive of neuron behaviour.

Fig. 10 shows that, for many neurons, the full model generalizes well to out of sample (unobserved) data, when compared to the AR model. This shows that inter-neuron connectivity is predictive of spiking rates.

Long range prediction is also possible using the recovered model parameters 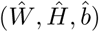 and the visual stimuli sequence to be presented ({*I*_*k*_(*t*)}). Instead of using the observed past spike trains *X*_*j*_(*t*) as regressors, we use the expected spiking rate *λ*_*j*_(*t*). This experiment iteratively computes the expected spiking rate for each neuron in the network using a fully observed external stimulation sequence and the past computed expected spiking rate. It is important to note that here we are computing the entire behaviour of the system given a stimuli sequence.

Formally, we define the long range spiking rate as

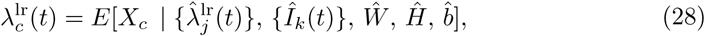

where 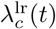 is the expected spiking rate at time *t* for neuron *c*,

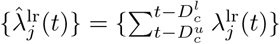 is the set of past expected spiking rates as seen through the seen through the boxcar influence function. boxcar influence function, and 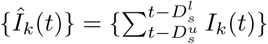 is the applied visual stimuli as seen through the boxcar influence function.

Here we compare the difference between the log likelihood when simulating the 2000 test samples with the stimuli that were presented versus a simulation of the system under a random equally likely stimuli selection.

The log likelihood of the sequences is computed against the observed spike trains for each neuron, experiments are repeated over 20 trials to provide error estimates. Results are shown in Fig. 11.

**Fig 11.**
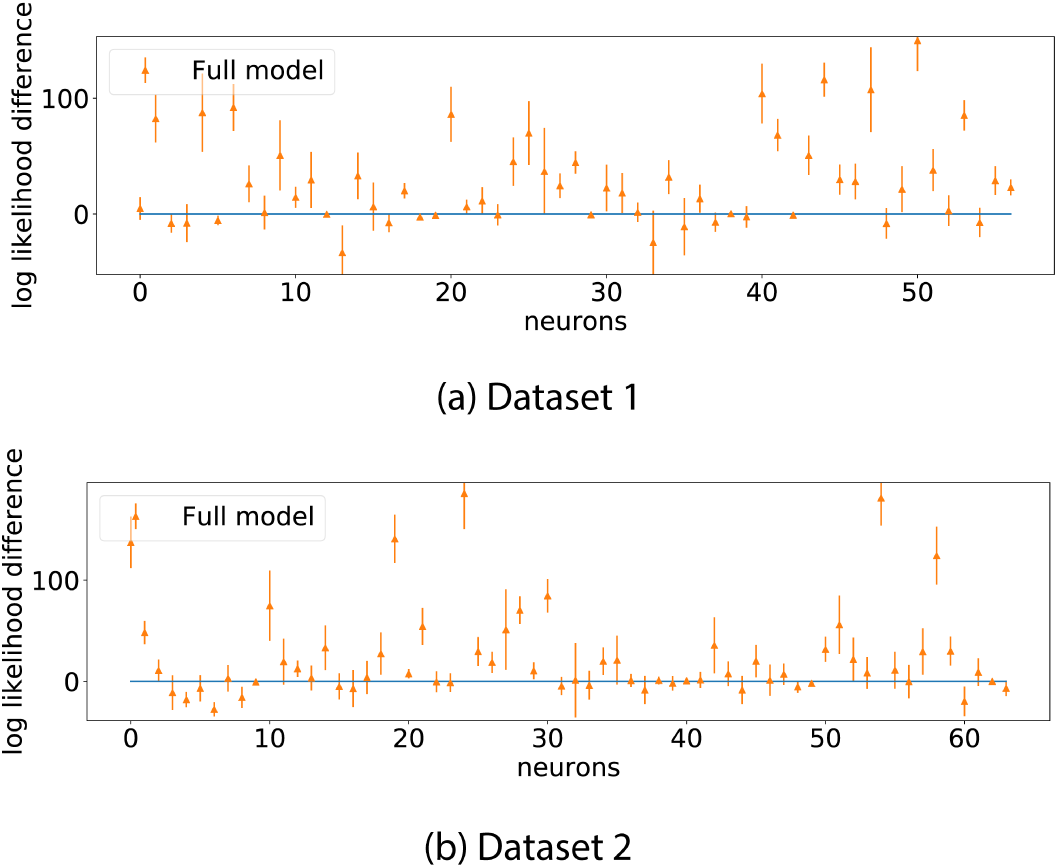
Difference in log likelihood of forecasted sequences, real stimulation sequence versus randomized stimulation sequences. Error bars represent one standard deviation over 20 trials.

Iterated simulations like these integrate the influence of stimuli to neuron connectivity and neuron to neuron connectivity. The large majority of neurons for both datasets showed a likelihood increase when presented with the true stimulation sequence, showing that the recovered models really capture interactions between regressors and spiking rates.

### Active learning on real data

In lieu of validating the proposed active learning framework on live animals, we utilize the recovered full model networks from Fig. 9 as ground truth. For this experiment, we simulate data using the recovered inter-neuron and stimuli response matrices (W, H) as ground truth. A separate model is trained on data drawn from this new simulation. We compare the performance of the network inference algorithm when samples are drawn uniformly from all 49 possible stimuli against samples drawn from the inferred active learning distributions.

Both strategies start from the same initial 1, 000 samples, and each intervention adds an additional 500 samples. As a reminder, the ground truth network was recovered from 9, 000 samples. At the beginning of each intervention step, we compute the best network estimate so far using Algorithm 1, and show the performance in recovering the set of regressor edges *PA*_*c*_ using the *F*_1_, precision and recall metrics as defined in Eq. (24). At this stage, the active learning strategy recomputes the stimuli probability distribution to apply for the following samples. These results are shown in Fig. 12. Additionally, Fig. 13 shows the stimuli probability distribution (*P*^*l*+1^) obtained from Algorithm 1, averaged over all realizations.

**Fig 12.**
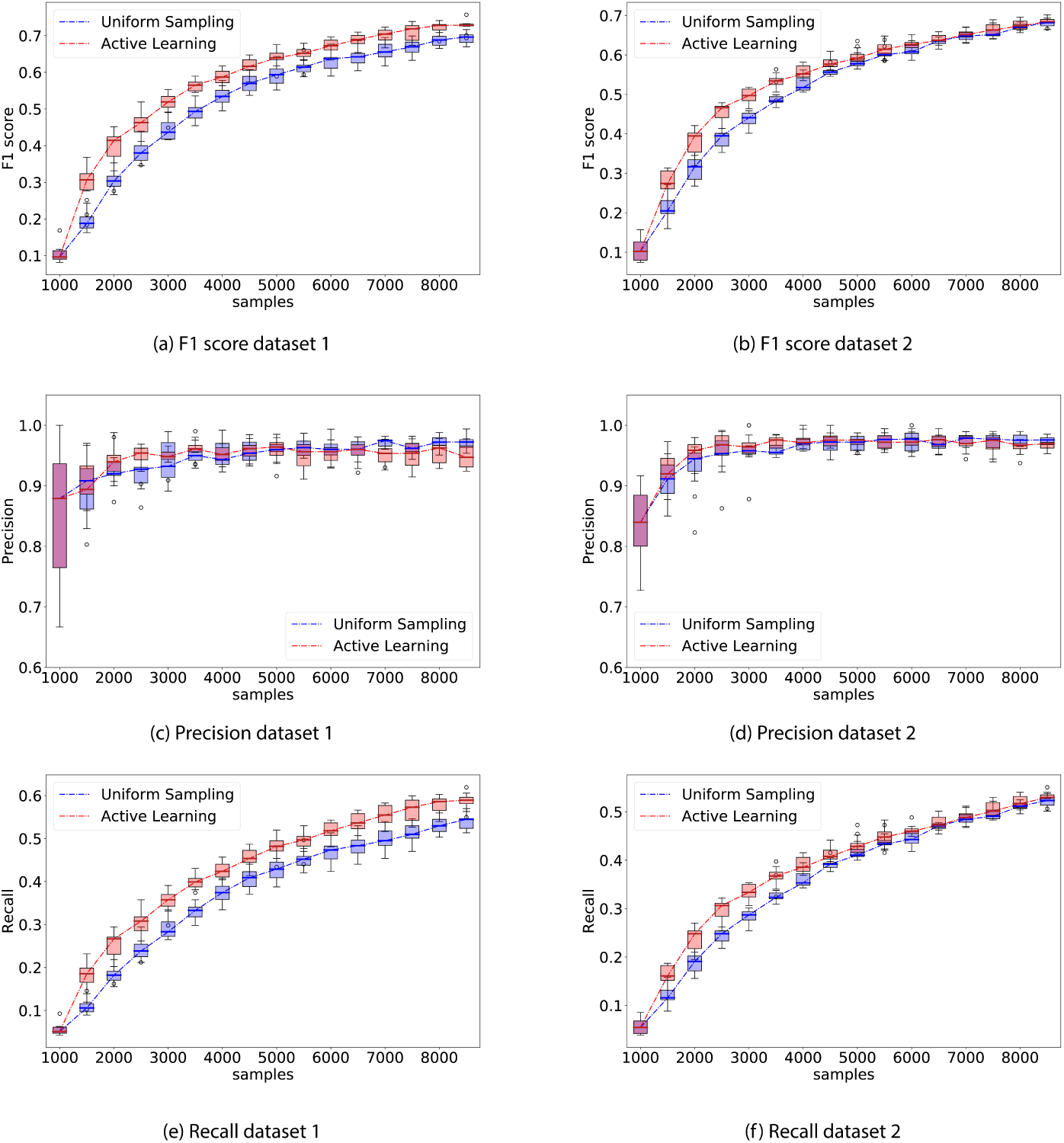
Comparison of performance between the proposed active learning (AL) method versus uniformly sampling from all stimuli. The experiment consisted of 500 sample interventions, with an initial 1, 000 sample observation. Whisker plot is obtained from 10 independent trials. Left column shows *F*_1_, precision, and recall performance on network recovered from dataset 1 and right column shows *F*_1_, precision, and recall performance on network dataset 2. On average, computing the Active learning intervention step took 108*s* ± 60*s* on dataset 1, and 86*s* ± 46*s* on dataset 2.

**Fig 13.**
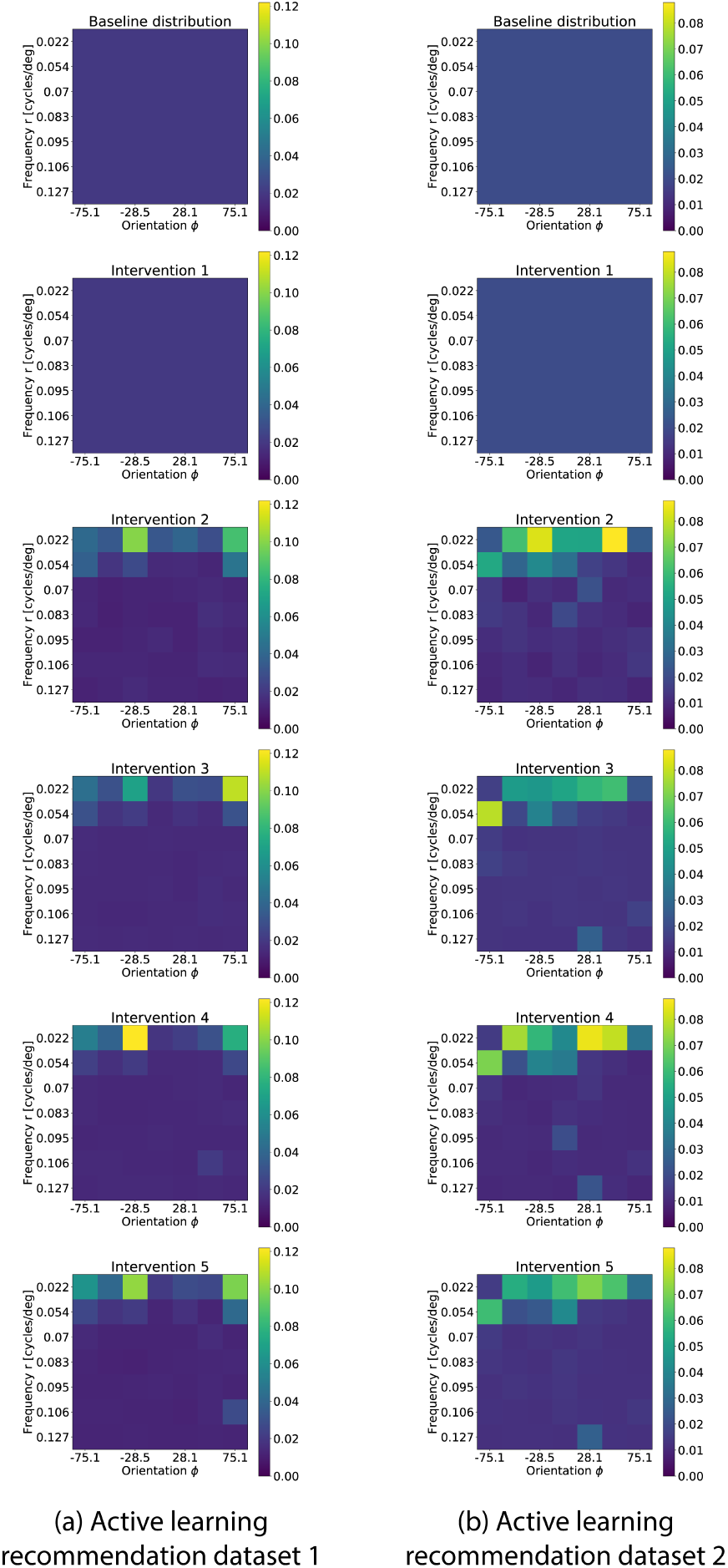
Distribution of recommended stimuli (*P*^*l*+1^) averaged across all realizations as a function of the number of interventions. Initially, the distribution is uniform (top). The experiment consisted of 500 sample interventions, with an initial 1, 000 sample observation. Left column shows the stimuli probability distribution history for dataset 1, while right column shows the distribution for dataset 2.

The *F*_1_ score of the active learning experiment is consistently better than random stimuli selection. While the performance gain is not large for these networks, the result spreads are tight and consistent; there is thereby no reason for not using the active learning strategy over random stimulation. We also observe that the AL algorithm preferentially presents low frequency stimuli, even though no explicit variable in the AL algorithm distinguishes between low and high frequency stimuli. In the following section we will show this is a reasonable result, since we found that neurons in these datasets tend to respond considerably more to low frequency stimuli.

### Network analysis

Now that we showed the performance of the proposed methods we will present some observations over the recovered biological models.

We first analyze the obtained input response matrices *H* for both datasets. Fig. 14 shows the percentage of neurons in each dataset that directly respond to each stimulation pattern. We can conclude that low frequency stimulation patterns have an out-sized proportion of directly responding neurons when compared to higher frequency patterns.

**Fig 14.**
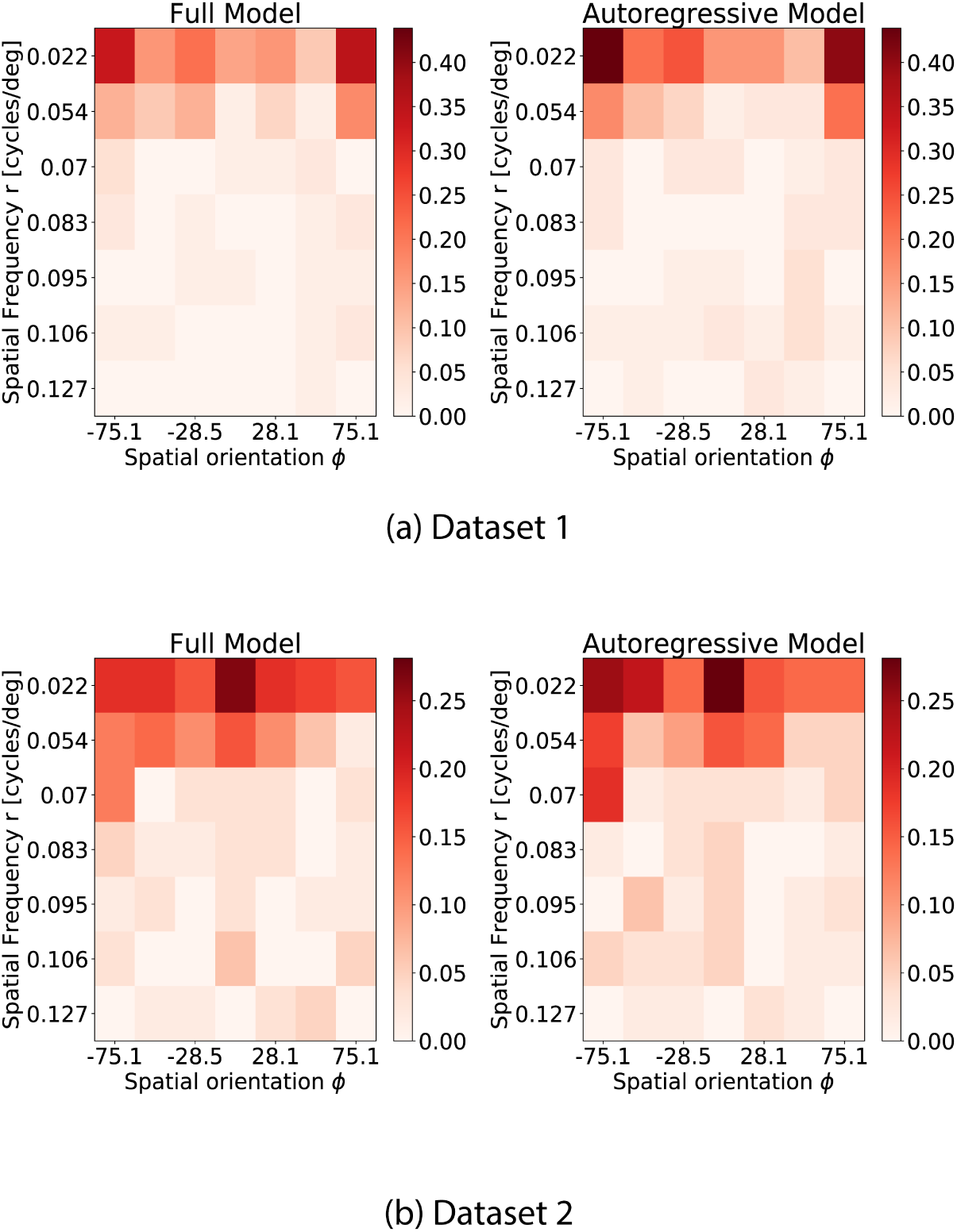
Percentage of neurons that have an excitatory or inhibitory response to each possible visual stimuli. Visual stimuli are represented in matrix form, where rows represent spatial frequency *r* and columns spatial orientation *Φ*. The value for each entry in the matrix is the percentage of neurons in the datasets that show a response to that visual stimuli. This visualization shows that both datasets show a large number of directly responding neurons for low spatial frequency visual stimuli.

For each dataset, we then examine the adjacency matrices *W* and extract two of the largest cliques in the network. Fig. 15 shows these cliques, and the spike trains of all neurons in the clique.

**Fig 15.**
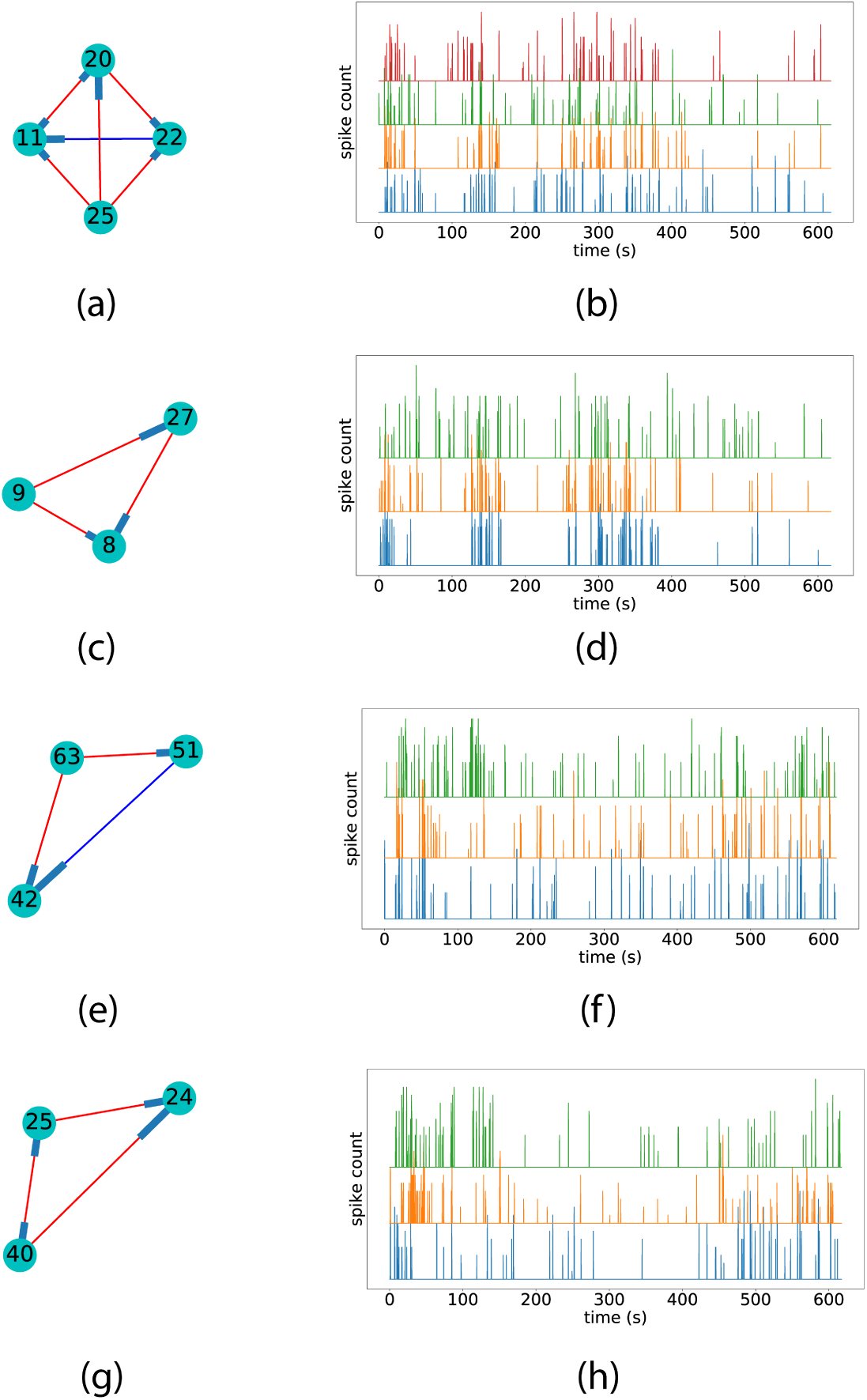
Figures a), c), e), and g) show the excitatory (red) and inhibitory (blue) edges detected for neuron cliques in datasets 2 (a) and c)) and 3 (e) and g)). Figures b), d), f), and h) show the spike time series of the neurons in the clique. The nodes are numbered according to the corresponding neuron index. We can visually see that spike trains from neurons in the selected cliques show similar spiking behaviour throughout the experiment.

We additionally count the occurrence rate of motif triplets in the recovered networks, and do a simple hypothesis test to check if these motifs could have arisen from a small-world network topology, Fig. 16 show these results. The high p-values observed in Fig. 16 show that the recovered network motifs are at least compatible with the small-world topology hypothesis. We also compared the distribution of number of children and parents per cell between the inferred networks and a small-world network ensemble. In Fig. 17 we can see that the inferred networks are compatible with a small-world topology in terms of number children and parents per neuron. Finally, we count the percentage of cells involved in multiple cliques, and further show the percentage of cells involved in multiple cliques of a set size. The percentage counts also appear to be within the expected counts of a small-world network, this is shown in Fig. 18.This is in accordance with other observations like the ones in [36–38].

**Fig 16.**
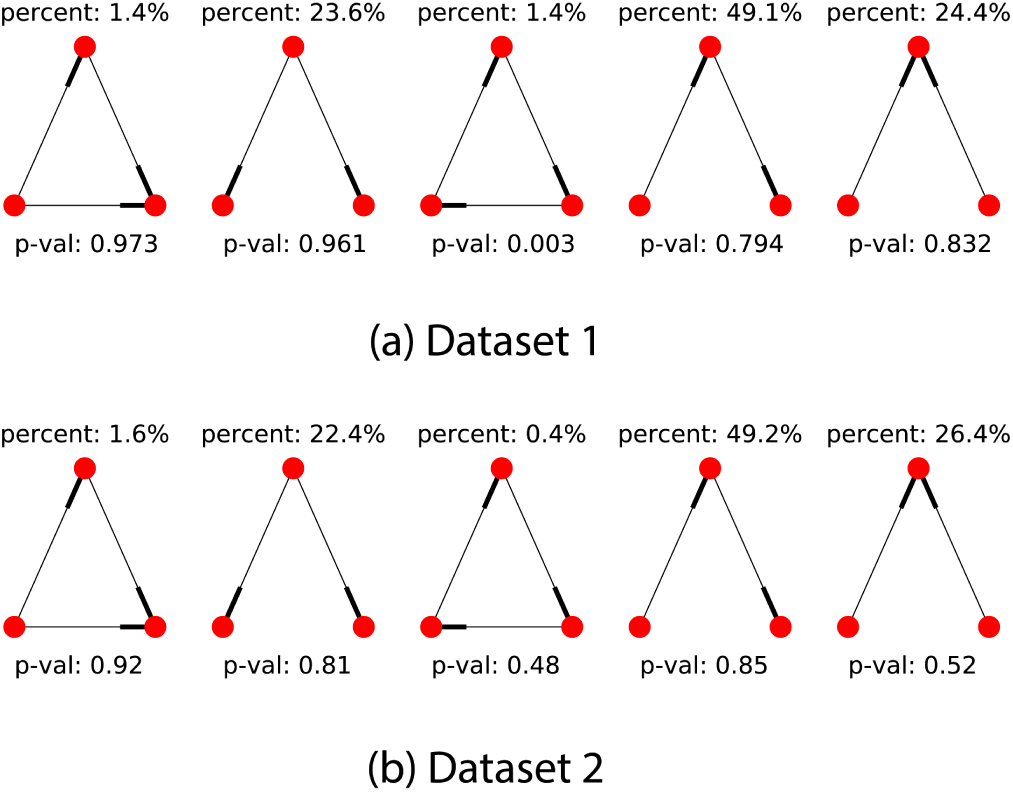
We count the occurrences of motif triplets for both datasets (we ignore edge weight and sign) by enumerating all neuron triplet combinations in the recovered networks and checking for graph isomorphism against all 5 motif triplet types. Top and bottom rows show results for datasets 1 and 2 respectively. We compare the obtained motif counts against a base model of small-world network topology and show the obtained p-values. These p-values are obtained by computation of the mean and standard deviation of each motif type in a small-world network with the same node count and edge density. The relatively large p-values obtained show that the small-world model is a good fit for the recovered network topology.

**Fig 17.**
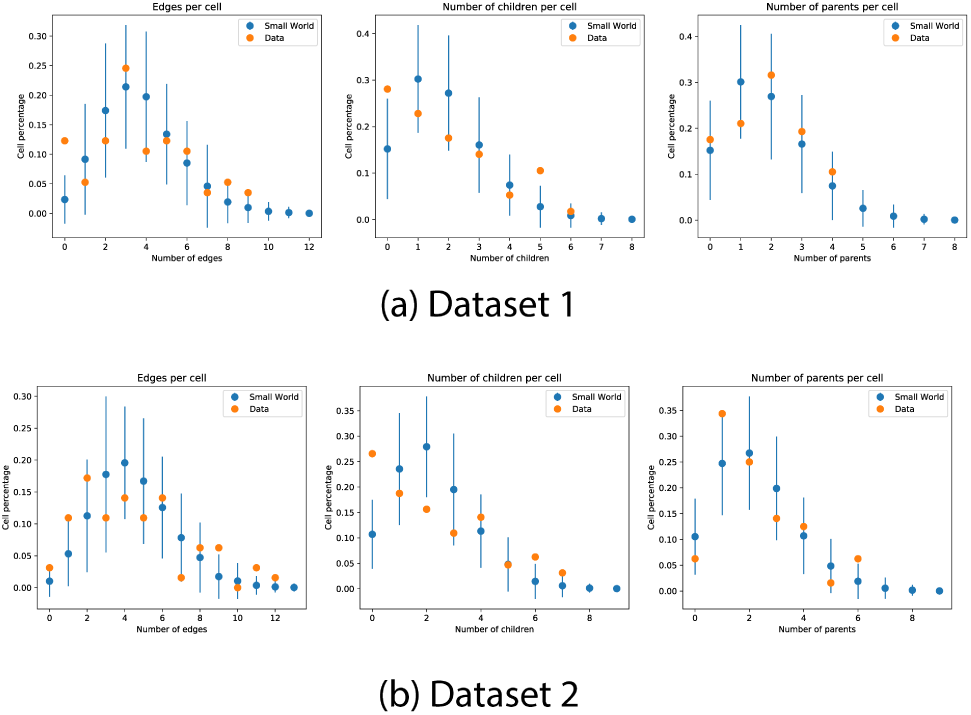
We compare the edge density distribution of the recovered inter neuron connectivity matrices in both datasets. Edge counts shown from left to right are all edges (number of neurons connected to node, either as parent or child), outbound edges (number of child nodes), and inbound nodes (number of parent nodes). The edge counts are compared against a base model of small-world network topology, error bars denote two standard deviations obtained from simulation of small-world networks with the same number of nodes and connectivity degree.

**Fig 18.**
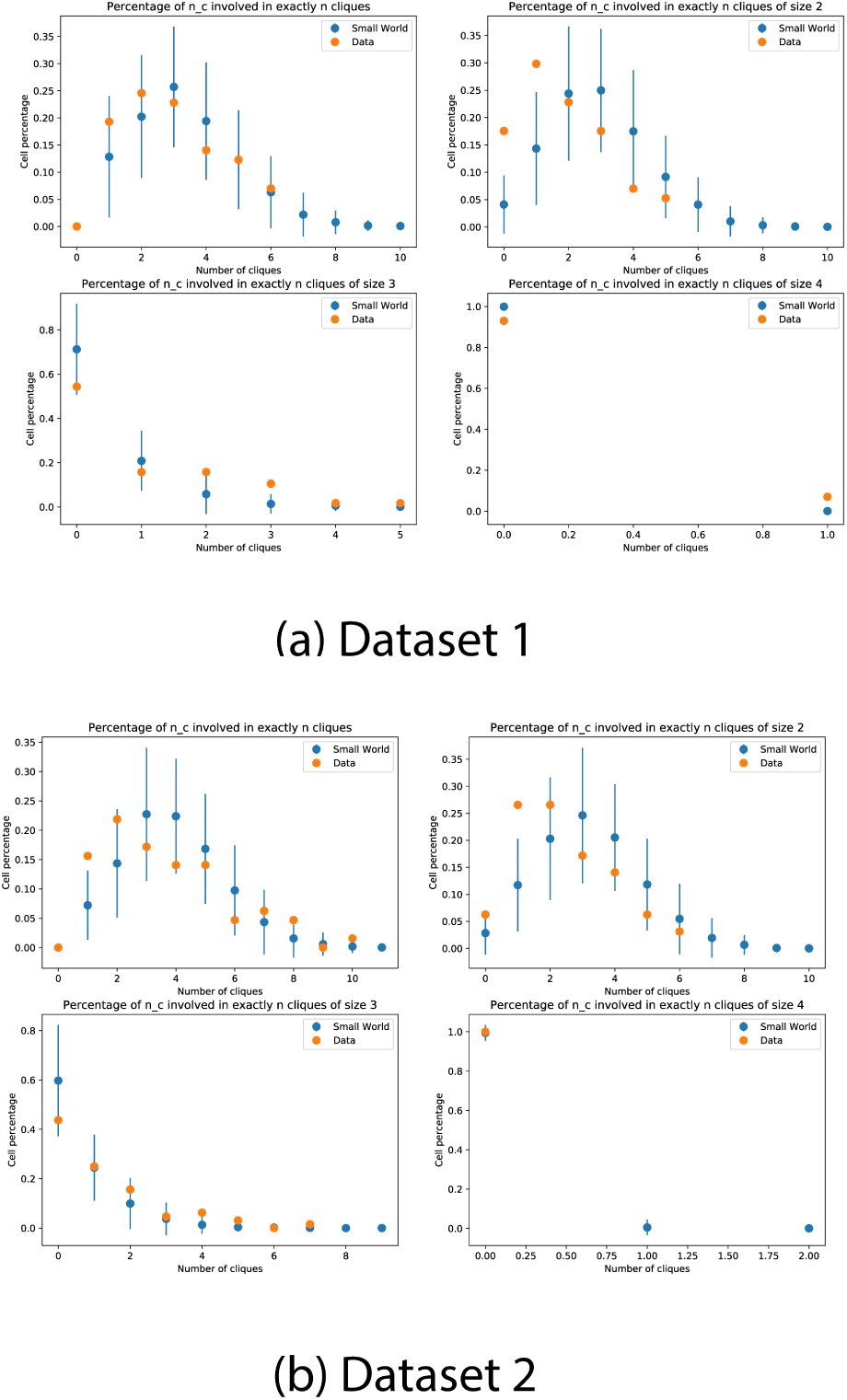
We count the percentage of cells participating in multiple cliques. The counts are compared against a base model of small-world network topology, error bars denote two standard deviations obtained from simulation of small-world networks with the same number of nodes and connectivity degree.

## Discussion

There is a large body of work done in reverse-engineering neural circuits from data across different modalities [40, 41]. Some of these methods learn a directed graph, which provides a compact and interpretable way of encoding Granger causal [42] relations between neurons and covariates. Several works have been published on the use of Gaussian Bayesian Networks to learn connectivity strengths from data [43, 44]. For cases where the framework is not applicable, as in the case of spike train time series, Generalized Linear Models (GLMs) have been successfully used [24, 45–51].

Parameters for these models are most commonly estimated using Maximum Likelihood Estimation or Maximum a Posteriori estimation [24, 46–51].The parameters obtained from the GLM can be interpreted as a directed graph capturing dependencies between variables of interest, or nodes (neuron spiking rates and visual stimuli). The presence of an edge in this graph represents a directed influence from one node to another, and the weight of the edge represents the magnitude of that influence. The graph can be made sparse with the use of subset selection, where only a limited number of edges are assigned non-zero weights (no influence). Subset selection can be performed using deviance tests [52] or by the use of priors [50].

In parallel, there is a corresponding push for actively estimating the best stimuli subset for network inference, with variants based on mutual information and Gaussian approximations of MAP parameters [24, 46, 48, 49, 53–62]

Methods like the one proposed in [53] use of mutual information for intervention selection, but rely on a specific Gaussian Bayesian Network framework, making them unsuitable for count data. Methods like [63] generalize D-Optimal factorial design for GLMs and to multi-level regressor covariates, but require full control over the regressor covariates.

The proposed variable selection algorithm is focused on subset selection of parent regressors. It is posed as an optimization problem where the objective is to find a set of regressors that minimize the BIC score, subject to a confidence interval restriction. Using BIC as the score to optimize fosters prediction improvement while penalizing model complexity. The use of p-value as a restriction criteria ensures that the regressors in the model are highly significant. A local minimal set of regressors is constructively obtained following a simple rule-set.

For the purposes of this work, the delay time window (boxcar length) was set directly to incorporate a delay of 5 time bins (∼ 0.3*s*). In a more general setting, the delay parameter could be set by fitting a model to data with various values of this parameter, and choosing the one with the highest likelihood.

The variable selection method was tested on simulated data and compared against oracle Lasso. It performed worse than oracle lasso for very small sample sizes, but otherwise proved to be better on the *F*_1_ and recall metrics. On settings where no ground truth is available, Lasso would require some other sub-optimal method for parameter tuning. The p-value restriction parameter present in the elastic-forward model selection algorithm is easily interpretable as the desireable confidence level on the regressor parameters, this makes the parameter easy to set beforehand for any experiment

Note that the time complexity of the forward model selection scales as the product of the number of available regressors and true edges in the network. For the worst case scenario (fully connected network), this means the forward model selection strategy scales quadratically with the number of regressors. For sparse networks, this computational cost can be brought down significantly by preselecting regressors based on the approximate p-value computation shown in Eq. (47).

The proposed active learning algorithm follows a simple design philosophy, it looks for promising edges not added into the model so far, and increases the appearance frequency of stimuli that drive up the spiking rate of the parent nodes of these edges. To achieve this, it defines a score for each possible stimulus based on the previous learned model, it takes into account the spiking rate difference (impact) of presenting one stimulus more frequently than the others on every parent node in the system, and weighs it by the potential log-likelihood improvement of adding every edge associated with this parent node to the model. The log-likelihood value of an edge is tightly related with the BIC score the elastic-forward model selection attempts to optimize.

Active learning proved to be faster than random stimulation in recovering edges on the simulated networks, this was especially true for edges whose spiking rate could be greatly affected by changing the stimuli distribution (first order connections). On the simulations from networks recovered from real data, the performance of active learning was consistently better than random stimulation, and the measured metrics had a tighter spread. There is therefore no reason not to use active learning during data acquisition.

We measured three basic properties of the recovered inter-neuron connectivity networks from on real data: edge distribution, motif type distribution, and number of cliques per neuron. Counts were compared to simulations of small-world networks with an identical number of nodes and connectivity degree. All three properties measured fell well within the expected values for these types of networks, pointing at a small-world-like structure in the recovered inter-neuron connectivity networks.

All codes used for this work are available at https://github.com/MartinBertran/ActiveLearningCortical

## Conclusion

In this paper we propose a simple framework for actively learning network connectivity for GLMs by selecting external forcing actions. The algorithm has the advantage of making relatively few assumptions on the exact distribution of the model, and the amount of control the experimenter has over the regressor covariates.

The use of a greedy regressor selector using BIC and Wald testing allows for an easy identification of edges that seem beneficial for the model, but do not yet have a sufficient number of interaction samples to be included in the model. By utilizing external triggers to these interactions, the algorithm prioritizes interventions that provide information over uncertain edges.

The greedy regressor selector outperforms the oracle Lasso in identifying the proper regressor subsets in simulations for non-small sample sizes, even when accounting for the oracle selection of the *l*_1_ prior.

Even on the recovered real datasets, the use of the active learning algorithm proved to be beneficial as well. The algorithm is very quick at recovering directly connected edges, making its application in conjunction with optogenetics an interesting proposition.

Finally, we note the method is not restricted to one modality or domain, it can be applied in any situation where there is a family of possible actions available to probe the activity of a network, in both biological and artificial systems.

We note that for all measured properties, the recovered network structure on the real datasets was consistent with a small-world topology.

## Appendix

### Derivation of the observed Fisher information

Given a neuron *c* and the MLE estimators 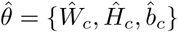, the element *k, j* of the observed Fisher information matrix 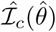 for the given model can be expressed as

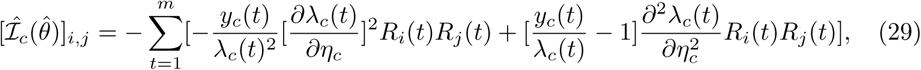

where *R* is the concatenation of all the considered regressors (neurons and stimuli),

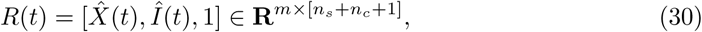

and the first and second derivatives of *λ*_*c*_(*t*) are,

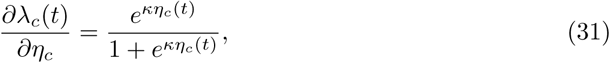

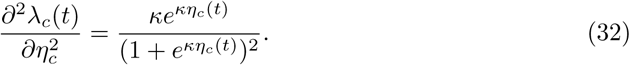

Replacing Eq. (31) and Eq. (32) in Eq. (29) we get:

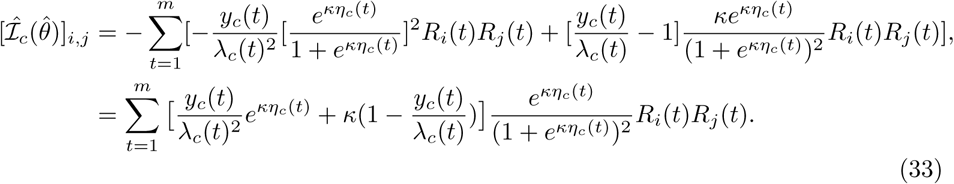

### Approximate Wald test dependence on model parameter *κ*

We now relate the Fisher information matrix to the *κ* parameter and other quantities that do not depend on the Maximum Likelihood estimate of the model. To achieve this, we make several approximations. We start by expressing Eq. (33)

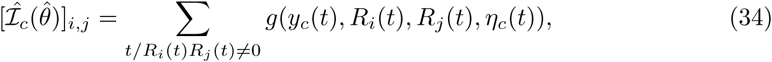

with

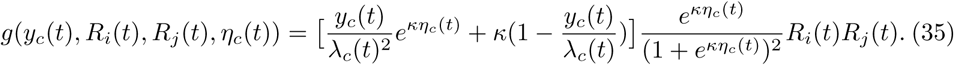

The first simplification we make is approximating the summation in Eq. (34) by its expected value. If we define *M*_*i,j*_ as the number of samples where *R*_*i*_*R*_*j*_ ≠ 0, we get

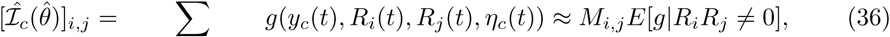

We further approximate this expression by moving the expectation inside the function,

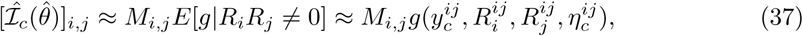

where

- 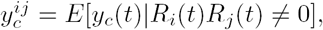
- 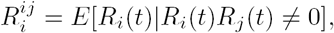
- 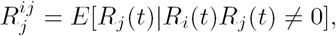
- 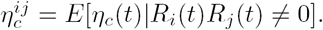

Going back to the full expression we get

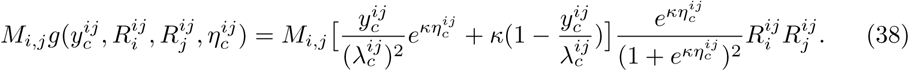

Notice that 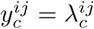 for a well fitted model, where 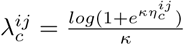 Then we have that:

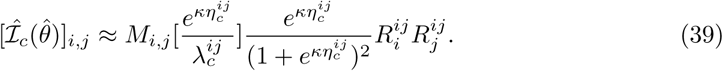

From the definition of *λ*_*c*_(*η*_*c*_) (Eq. (4)) we can express 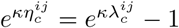 and obtain

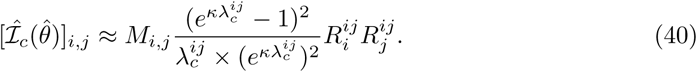

We relate this Fisher information matrix approximation to the z-score of any given regressor. To avoid computing the inverse of the Fisher information matrix, we take the following approximation:

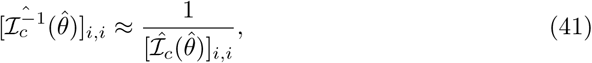

where

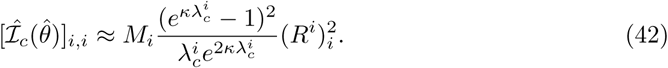

By replacing this approximation into Eq. (9), the Z-score of regressor *i* on neuron *c* can be very roughly approximated to:

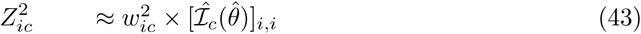

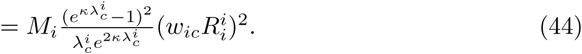

Our final task is relating the term 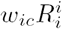 to quantities that do not depend on the Maximum Likelihood estimate. To do this, we again note that for any well fitted model we can write:

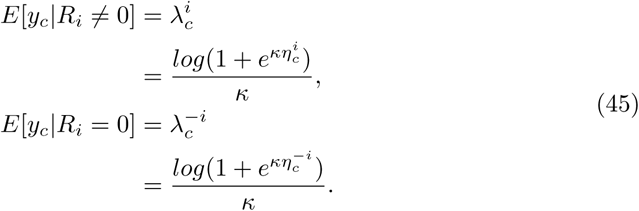

We conveniently express 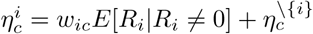 where 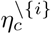 is the contribution of all regressors beside regressor *i*. Similarly, we express 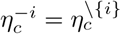, where we further assumed that the total contribution of all other regressors to the *η*_*c*_ parameter is mostly independent on the state of regressor *R*_*i*_.

Working both equations, we obtain:

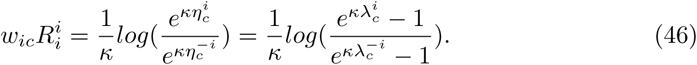

We finally conclude that, after several approximations, the Z-score of a parameter can be roughly approximated with an expression that only depends on the rates of our data and the parameter *κ* of our model,

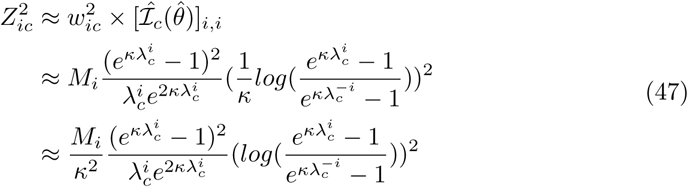

Using this approximation, we can gain some intuition on how the Z-score will behave under certain conditions. In particular, we see that the Z-score depends approximately linearly on the number of observed interactions, it also depends on what is essentially a ratio between *E*[*y*_*c*_*|R*_*i*_ ≠ 0] and *E*[*y*_*c*_*|R*_*i*_ = 0]. The dependence on the *κ* parameter is not easy to discern at a glance, but we can observe how the Z-score behaves for various combinations of spiking rate parameters and *κ* values.

Fig. 19 shows how this approximate Z-score function behaves for several *κ* as a function of the fractional rate 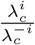 and 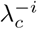. Rates and base spiking rates where selected to be in accordance to the rates measured in our real datasets.

**Fig 19.**
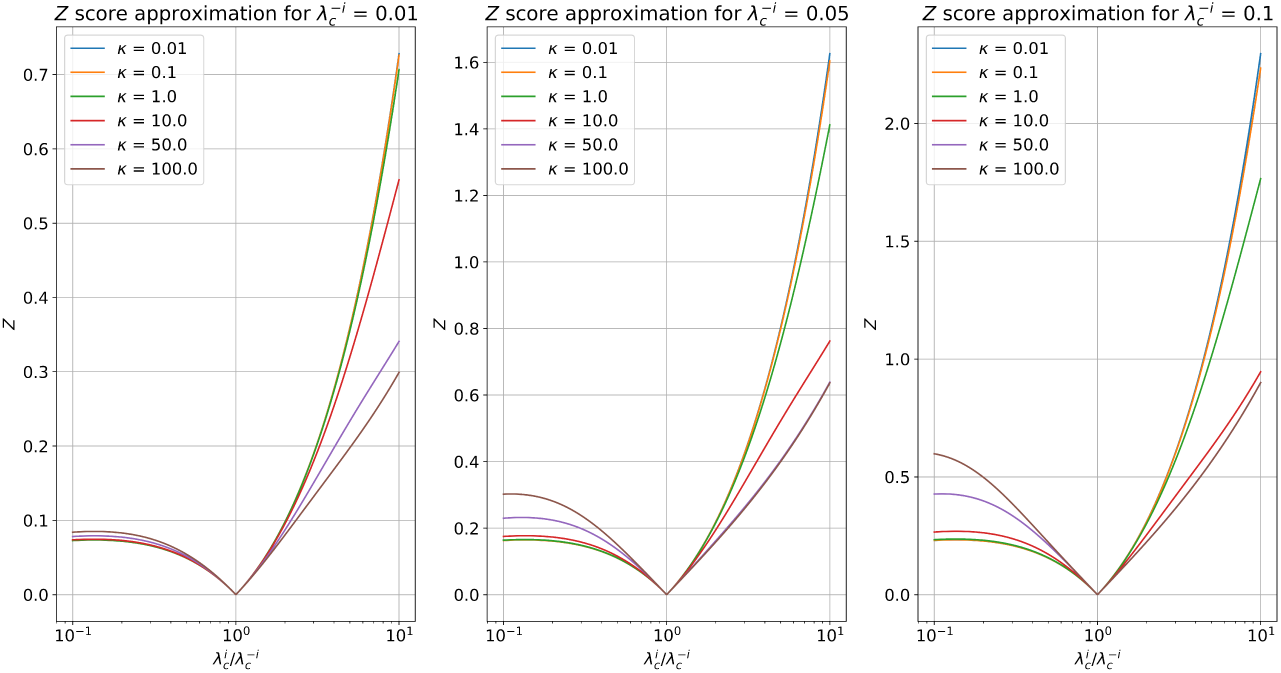
Single sample (*M* = 1) Z-score approximation according to Eq. (46) for several *κ* parameters as a function of the ratio 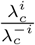 and 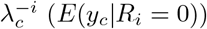. The 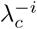 values were chosen to be representative of the observed rates in our real datasets.

Numerical issues arise from picking very large values of *κ*. We decided to choose *κ* = 10 for all experimental results since it seemed to provide a nice middle ground between dampening the Z-score for low spiking rate inhibitions and not giving disproportionately large Z-scores for small excitatory rates.

### Fisher information: Model selection and the under-sampled regime

We analyze the behaviour of ill-conditioned but non-singular Fisher matrices, where there is a potentially large difference between the largest and smallest eigenvalues present in the Fisher information matrix, and show how the Wald test present in the model selection strategy copes with this issue. We use this analysis as we take the limit of the smallest eigenvalue going to 0 to illustrate what this test does for singular matrices.

Overall, we provide a link between the smallest eigenvalue of the Fisher information matrix, and the largest observed parameter variance we obtain from the diagonal elements of the inverse Fisher information matrix. From this we finally conclude that regressor subsets that produce nearly singular matrices are consistently rejected from our model selection process.

We first note that, for any non-singular matrix *A*, if *λ* is an eigenvalue of *A*, then *λ*^-1^ is an eigenvalue of *A*^-1^. So the inverse of a matrix with a very small eigenvalue has a correspondingly large eigenvalue.

We also note that the Fisher information matrix is positive (semi) definite, we can thus sort its eigenvalues in descending order and write *λ*_1_ *≥ λ*_2_ *≥…≥λ*_*r*_ *≥*0, where *r* is the number of rows (regressors) of the matrix. The corresponding eigenvalues of the inverse Fisher matrix are ordered: 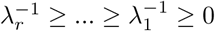

Since the inverse Fisher information is symmetric, we can use the Schur-Horn theorem to see that its diagonal elements are majorized by its eigenvalues. By definition of majorization, this means that:

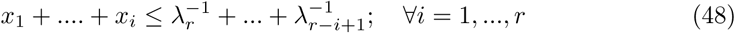

where *x*_*i*_ is the i-th largest entry in the diagonal elements in the inverse Fisher matrix, the equality holds exactly for *i* = *r*. These values are the fisher variances of the regressors in Eq. (9), sorted by descending variance.

From this, we can easily see that the largest observed fisher variance is at least

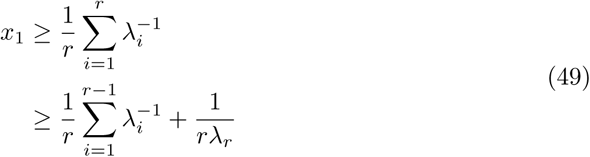

From this we can conclude that as the smallest value of the Fisher information matrix goes to 0, at least one regressor’s Fisher variance grows arbitrarily large. Relating this back into Eq. (9) and Eq. (12) we can see that for any reasonable p-value threshold, a parameter subset *PA*_*c*_ yielding an ill-conditioned Fisher information matrix will be rejected.

### Regressor selection: fraction parameter *ν*

We evaluate the effect of varying the sample fraction *ν* used for the random subset selection step of the elastic-forward model selection method. The *ν* parameter is tested in the 0.5 to 0.9 ranges. This is compared against the baseline oracle lasso method and model selection where no random subset selection is performed (no-subset).

All methods are evaluated on samples drawn from simulated network SW1CL using the *F*_1_, precision, and recall performance metrics. Results are shown in Fig. 20.

**Fig 20.**
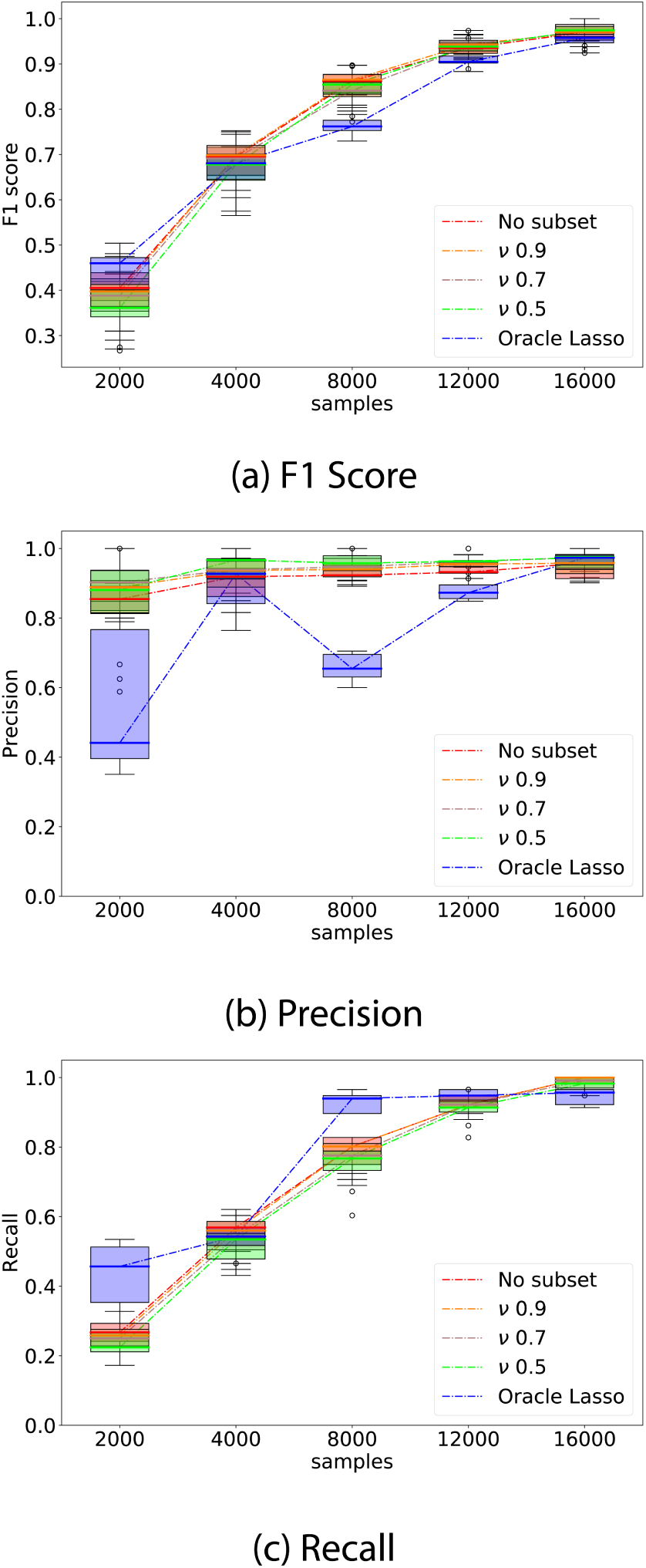
Whisker plot of performance indicators as a function of number of samples; elastic-forward BIC selection is compared using several *ν* parameters against both oracle Lasso and the special case where the whole dataset is used at once, without splitting into random subsets (no-subset). The plots show relatively little difference between the various *ν* parameters, but the use of the *ν* parameter is better performing overall to both oracle lasso and no-subset model selection.

We can observe from Fig. 20 that all *ν* parameters perform similarly for all but the smallest sample sizes. Additionally, the use of no-subset model selection had a large performance decrease for small sample sizes when compared to all *ν* parameters, but similar performance on larger sample sizes.

### Poisson model selection on non-Poisson models

A natural question that may arise is if the Poisson GLM model can still capture directed interactions between neurons when the data does not come from a Poisson distribution.

To that effect, we repeated the simulated experiments using the same connectivity matrices defined for networks SW1CL and SW3CL, but this time, the spiking activity was obtained using a Leaky Integrate and Fire model [64].

The equations of the model can be summarized as

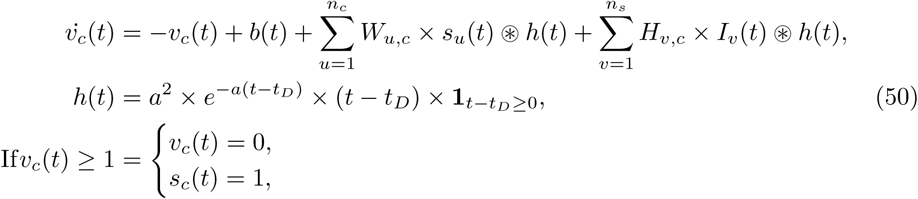

where *v*_*c*_ is the membrane potential of neuron *c*, and *s*_*c*_ its corresponding spike train.

*W* and *H* are the inter-neuron and stimuli-neuron connectivity matrices respectively. The parameter *h*(*t*) represent the influence kernel, and depends on the synaptic density (*a*) and kernel delay (*t*_*D*_). *b*(*t*) is the direct current parameter.

For our simulations we sampled *b*(*t*) from a uniform distribution (*b*(*t*) ∼ *U* [0*, b*]), where *b* was chosen such that the average spiking rate of each neuron is the same as in the original Poisson GLM simulations. The *W* and *H* connectivity matrices were linearly scaled with respect to the original SW1CL network to preserve the conditional spiking rates. For these simulations, we set *a* = 1.5 and *t* _*D*_ = 2.

Fig. 21 compares the performance of the Active Learning algorithm and Uniform sampling when applied to the non-Poisson, Leaky Integrate and Fire network.

**Fig 21.**
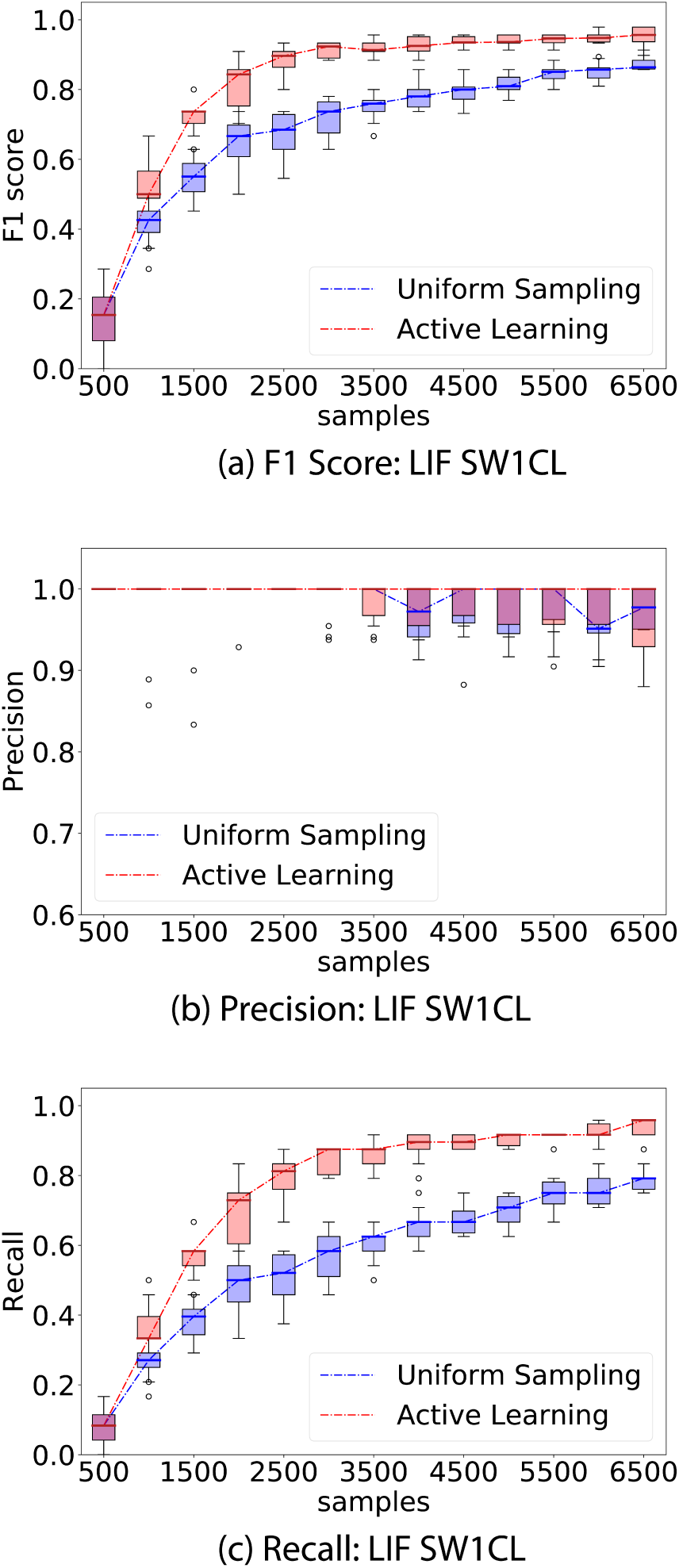
Comparison of performance between the proposed active learning method versus uniformly sampling from all stimuli. Spiking trains were simulated using a Leaky Integrate and Fire model, but the recovered networks were done using the proposed Poisson GLM. The experiment consisted of 500 sample interventions, with an initial 500 sample observation. Whisker plots are obtained from 10 independent trials.

From Fig. 21, we see that, at least for this tested configuration, the Poisson GLM is still able to recover the correct directed connections even when the originating data model does not follow a Poisson distribution.

### Table of defined variables

**Table 1.**
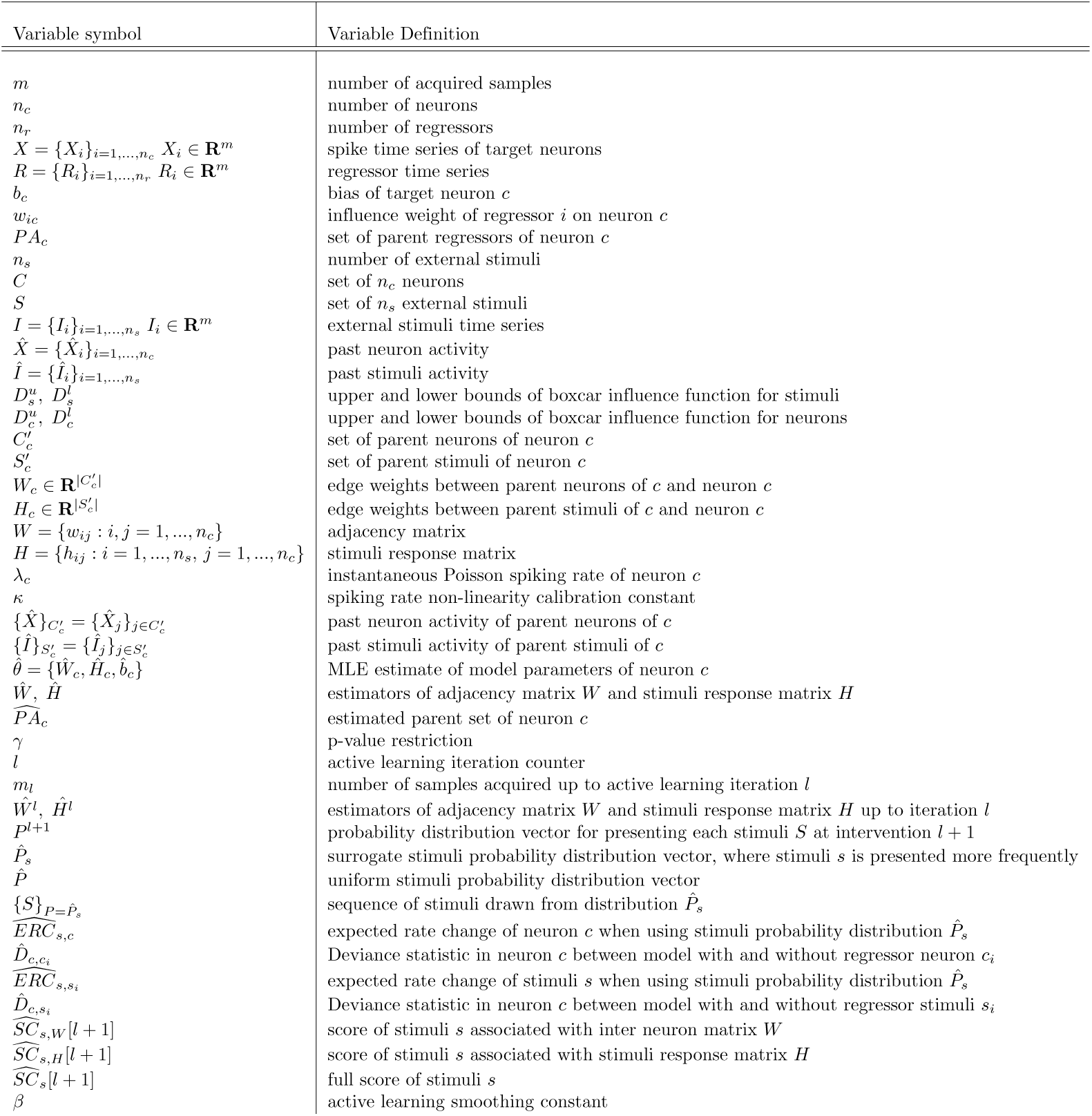
Summary of defined variables

## Algorithms

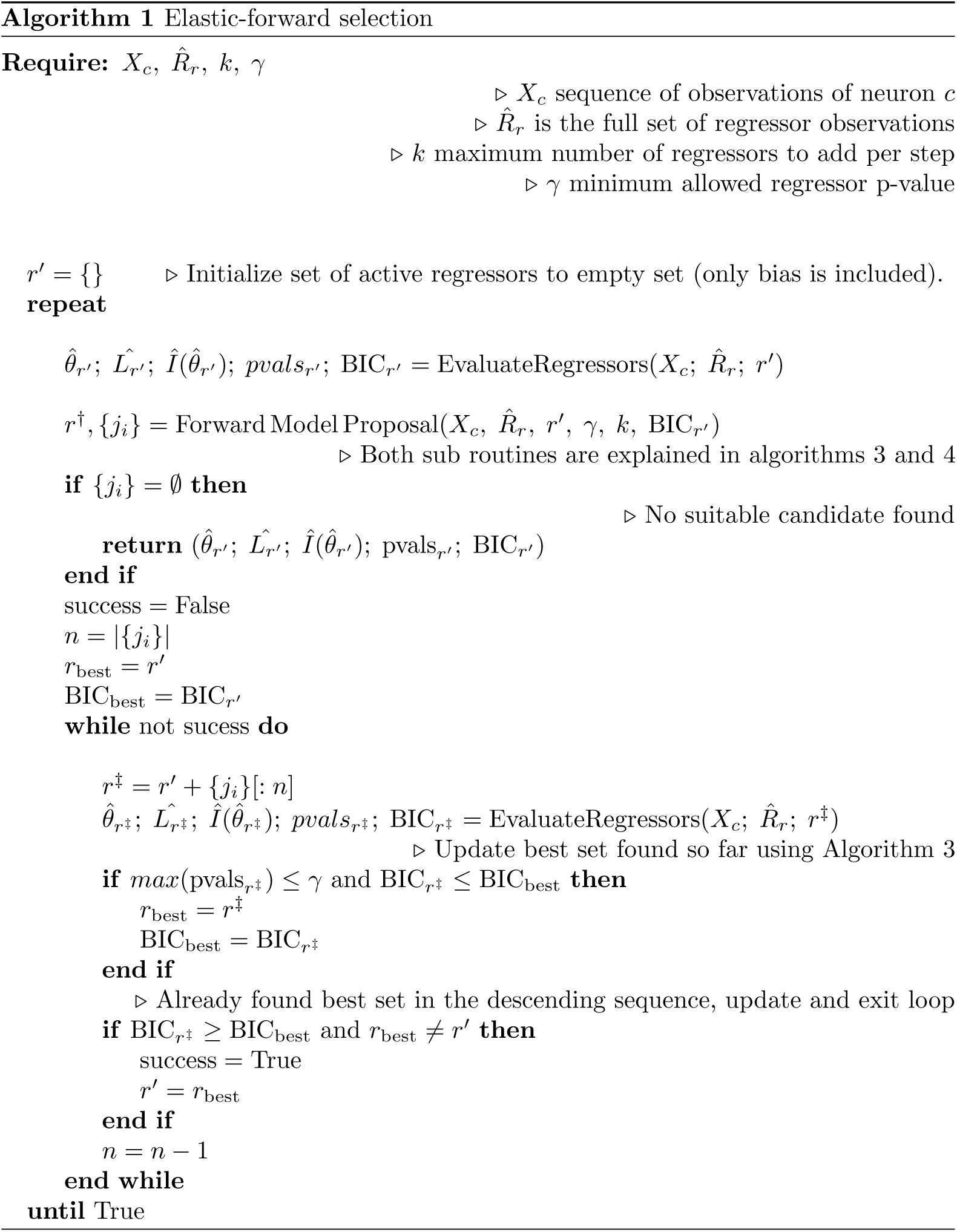

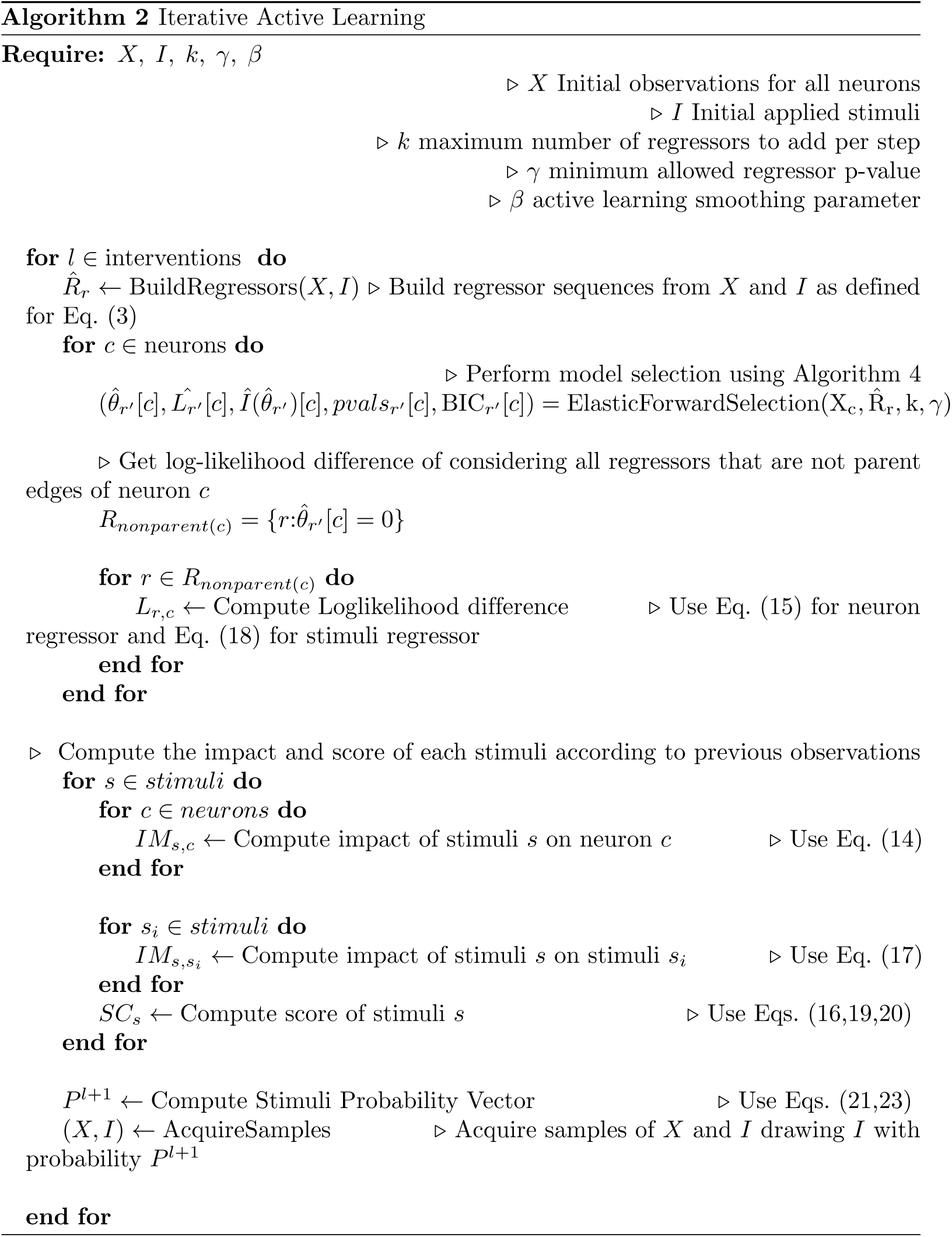

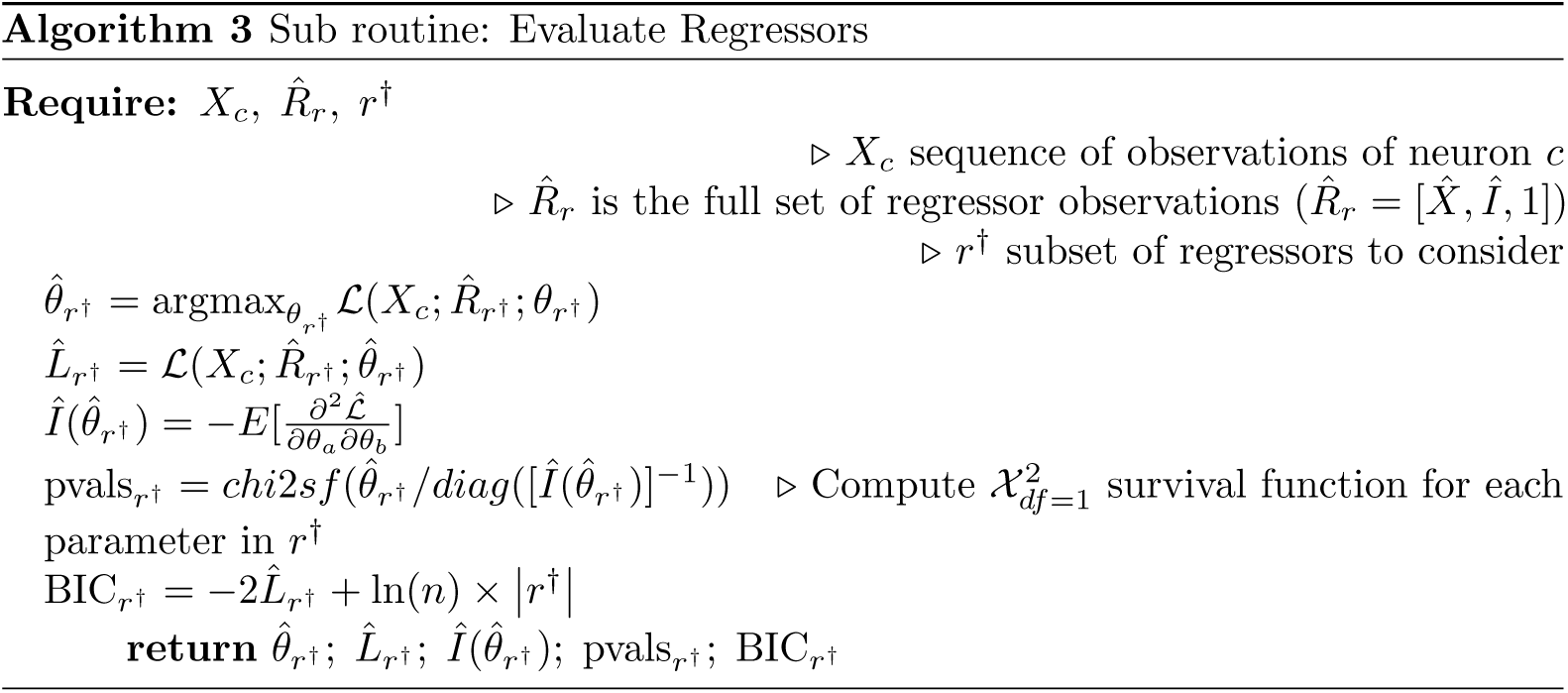

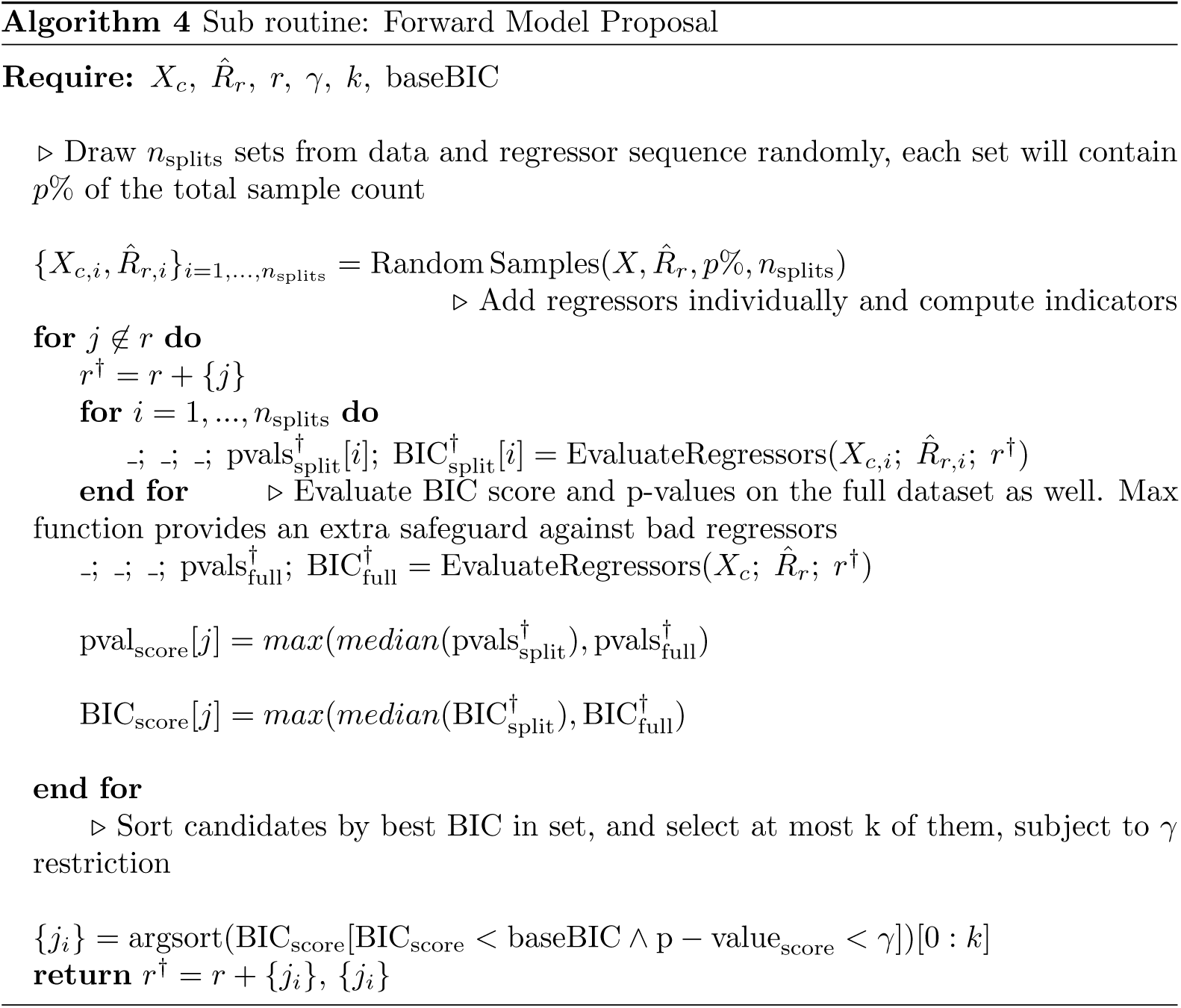

## Acknowledgments

We thank Prof. Gregory Randall for important discussions during the early stages of this work. This work has been supported by NIH, NSF, and DoD.

